# ST6GAL1-mediated sialyl linkage switching in tumor-associated macrophages drives cancer-promoting nanotubes carrying α2,6-sialylation in anti-inflammatory cells

**DOI:** 10.1101/2024.11.28.625934

**Authors:** Priya Dipta, Naaz Bansal, Zeynep Sumer-Bayraktar, Arthur Chien, Julian Ugonotti, The Huong Chau, Merrina Anugraham, Daniel Kolarich, Boaz Tirosh, Rebeca Kawahara, Morten Thaysen-Andersen

## Abstract

Tumor-associated macrophages (TAMs) form functionally diverse populations of innate immune cells in the tumor microenvironment (TME). Pro- and anti-inflammatory TAMs are central to cancer progression by shaping inflammation and immune (im)balance, but it remains unknown if polarization-induced remodeling of the TAM glycocalyx critical for cellular communication occurs within the TME. Taking a systems glycobiology approach, we here firstly used cell surface-focused glycomics and lectin flow cytometry of *ex vivo* polarized monocyte-derived macrophages to demonstrate profound sialyl linkage switching of the surface *N*-glycome in pro-inflammatory (α2,3-sialo-favored) and anti-inflammatory (α2,6-sialo-dominant) macrophages. In contrast, no polarization-induced alterations in sialylation were observed in the surface *O*-glycome. ST6GAL1 that modifies *N*-glycans with α2,6-sialylation was elevated in anti-inflammatory compared to levels in pro-inflammatory macrophages providing a mechanistic basis for the sialyl linkage switching, which was supported by *ST6GAL1* silencing. Interestingly, SNA-focused lectin cytochemistry of anti-inflammatory macrophages revealed dense networks of dynamic α2,6-sialylated protein-based nanotubules forming inter-connecting cellular structures that were absent in pro-inflammatory macrophages. Temporal *ST6GAL1* silencing in anti-inflammatory macrophages caused nanotubule disintegration as evidenced by SNA and biotin fluorescence microscopy. Moreover, live cell recordings of anti-inflammatory macrophages cultured with and without colorectal cancer (CRC) cells showed reduced macrophage motility, attenuated inter-macrophage and macrophage-CRC cell interactions and diminished CRC cell proliferation upon ST6GAL1 disruption indicating functional roles of the α2,6-sialylated nanotubules. Finally, sialyl linkage switching was recapitulated in pro- and anti-inflammatory TAMs from tumor tissues of patients with advanced CRC. We report on the mechanistic basis for and functional consequences of glycocalyx remodeling accompanying TAM polarization.

## 1. Introduction

Macrophages are long lived and plastic innate immune cells that contribute to the first line of immune defense in tissues by regulating inflammation and eliminating harmful agents including pathogens and cancer cells [1–3]. While macrophages predominantly arise from bone marrow-derived blood monocytes, tissue-resident and self-replenishing macrophages are now also recognized as important cell populations within the innate immune system [4, 5]. Forming a critical immune component of many, if not most, cancers, macrophages prevalently infiltrate and impact developing tumors and the proportion of tumor-associated macrophages (TAMs) in the cellularly diverse tumor micro-environment (TME) is reportedly as high as 30-50% [6–8].

Upon infiltration into tumors, TAMs rapidly undergo polarization into discrete macrophage populations displaying distinct functional programs in response to chemical cues within the TME [9–11]. In the early tumor stages (environments rich in lipopolysaccharide [LPS], toll-like receptor [TLR] ligand, and IFN-γ), TAMs predominantly undergo classical polarization to form pro-inflammatory macrophages that are regarded as cancer-suppressing cells [12, 13]. Due to their anti-tumor properties, pro-inflammatory macrophages delivered through intra-tumor injections have been explored in cancer therapy [12, 14]. In advanced tumor stages (featuring IL-4 and IL-13), TAMs undergo alternative polarization to form anti-inflammatory macrophages regarded as a cancer-supporting phenotype [15–18]. Enabled by their extracellular matrix (ECM) remodeling capabilities, anti-inflammatory macrophages promote tumor growth [19, 20], support cancer stem cells and induce humoral immunity [21, 22]. Consistent with such tumor-promoting traits, studies have shown associations between the density of anti-inflammatory macrophages and poor prognosis of patients with colorectal cancer (CRC) [23], hepatocellular carcinoma [24] and non-small-cell lung carcinoma [25, 26].

Glycosylation is a functional post-translational modification (PTM) of proteins essential for cellular and tissue functions and homeostasis [27]. By impacting a broad range of cellular processes including cell differentiation, adhesion, migration, and defense, glycosylation is central to maintain immune balance and regulate inflammation [28, 29]. Being a dynamic PTM exquisitely sensitive to the local cellular environment, diverse glycosylation features have repeatedly been linked to the immune imbalance underlying cancer [30, 31]. Most prominently, aberrant sialylation involving the under- or over-expression of sialo-glycans capped with *N*-acetylneuraminic acid (NeuAc) residues or an altered sialyl linkage type distribution (e.g. α2,3- or α2,6-linked sialylation) are increasingly regarded as molecular hallmarks across various cancer types [32, 33]. Cancer-associated aberrations in the sialylation patterns are mediated by sialyltransferases (STs), a broad class of anabolic glyco-enzymes that undergo expression changes in the TME [34–36]. Enabled by their exposed (terminal) position on glycoproteins embedded into the cellular glycocalyx, altered sialylation of cancer cells contributes to immune evasion by generating damage-associated molecular patterns (DAMPs). These “don’t eat me” signals are recognized by sialic acid-binding immunoglobulin-like lectins (Siglecs) on tumor-infiltrating immune cells. Changes to the interactions between Siglecs and sialoglycans (Siglec ligands) and may serve to dampen the immune response in favor of tumor progression [37, 38].

The glycocalyx of macrophages is rich in sialylated *N*-glycans as well as core 1- and 2-type sialo-*O*-glycans [39–43]. We have longitudinally surveyed glycosylation during monocyte-to-macrophage transition and found that while the cellular glycoproteome remains relatively constant during this metamorphosis, prominent glycocalyx remodeling of particularly the sialo-glycoepitopes was observed during late-stage maturation to naïve (unpolarized) macrophages [43]. While we did not explore glycome remodeling and functional consequences in fully polarized (pro- and anti-inflammatory) macrophages, the study illustrates the dynamic glycophenotypic modulation that accompany myeloid cell transformation also mirrored in other innate immune cell populations [44].

Functionally, the glycocalyx is known to impact key macrophage processes including their responsiveness to and surveillance of cancer cells [30]. For example, a recent study found that genetic and enzymatic desialylation of tumor cells delay tumor growth by repolarizing TAMs towards a pro-inflammatory anti-tumor phenotype [45]. Although providing neither molecular details nor mechanistic insight supporting their observations, glycosylation on the surface of macrophages was also suggested to guide the polarization towards either pro- or anti-tumorigenic populations [46–48]. Despite this growing body of literature highlighting a functional link between glycosylation and the role of TAMs in cancer, less is known of the molecular changes accompanying macrophage polarization into pro- and anti-inflammatory phenotypes and how such glycophenotypic alterations of the TAMs may impact tumorigenic processes to manipulate and shape the outcome of developing cancers.

In this study, we employ system glycobiology approaches to define with precision and subcellular resolution the macrophage glycocalyx in paired pro- and anti-inflammatory macrophage populations robustly differentiated and polarized from donor monocytes. Prompted by the observations of profound polarization-induced sialyl linkage switching, which were recapitulated in TAMs present in tumor tissues from patients with advanced CRC, we then employed methods in molecular and cellular glycobiology to demonstrate that ST6GAL1 is the glyco-enzyme that drives the glycophenotypic differences between pro-inflammatory (α2,3-sialylation favored) and anti-inflammatory (α2,6-sialylation dominant) macrophages. Notably, lectin fluorescence microscopy revealed dense networks of α2,6-sialylated nanotubules in anti-inflammatory macrophages forming inter-connected cellular structures that disintegrated upon ST6GAL1 silencing, whereas pro-inflammatory macrophages lacked such nanotubular structures. Finally, live cell recordings were used to document that the ST6GAL1- mediated sialyl linkage switching affects the motility of anti-inflammatory macrophages, modulates macrophage-macrophage and macrophage-CRC cell interactions and alters CRC cell proliferation pointing to potential functions of the dynamic and structurally heterogenous α2,6-sialylated nanotubules. The study provides mechanistic insights into the glycobiology of TAMs, which, in turn, elevates our understanding of cellular communication within the TME and opens avenues for advanced immuno-therapies directed against cancer.

## 2. Materials and Methods

### Chemicals and reagents

Chemicals and reagents for the -omics experiments were of LC-MS grade and obtained from Sigma-Aldrich, Merck or Thermo-Fisher Scientific unless otherwise stated. Granulocyte-macrophage colony-stimulating factor (GM-CSF, #130093865), IL-4 (#130093917) and IL-13 (#130112409) were from Miltenyi Biotec, Sydney, Australia. Interferon-gamma (IFN-γ, #PHC4031), lipopolysaccharide (LPS, #00497603), anti-CD14 (#11014941), anti-CD68 (#11068941), anti-CD86 (#12086941, #51086941), and anti-CD206 (#53206941) monoclonal antibodies as well as streptavidin AF488 (#S32354), ActinGreen 488 ReadyProbe against F-actin (#R37110), and 4′,6-diamidino-2-phenylindole (DAPI, #H1398) were from Thermo-Fisher Scientific. Cy3-conjugated *Sambucus nigra* lectin (SNA, #21761140-2), and Cy3- and FITC-conjugated *Maackia amurensis* lectin (MAL-I, #21761120-1, #21761036-1) were from Bio-world, Ohio, USA.

### Isolation of PBMCs and monocytes

Buffy coat of healthy individuals was provided by Australian Red Cross Lifeblood (Sydney, Australia) with approval from Macquarie University Human Research Ethics Committee (#52021780424399). Donor details were deidentified. Peripheral blood mononuclear cells (PBMCs) were isolated from buffy coat by density using Lymphoprep (#07801, StemCell Technologies) with centrifugation at 1,200 x *g* for 20 min at room temperature. The intermediate buffy layer was collected and washed with PBS containing 2 mM EDTA and 1% (w/v) bovine serum albumin (BSA). CD14+ monocytes were isolated through negative selection using a Pan monocyte isolation kit (#130096537, Miltenyi Biotec) as described [49]. Isolated CD14+ monocytes were validated for purity (≥95%) by flow cytometry (**Supplementary Figure S1A-B**).

### Differentiation and polarization of macrophages

Monocytes were washed and resuspended in Roswell Park Memorial Institute media 1640 (RPMI, #11875093, Gibco) supplemented with 10% (w/v) heat-inactivated fetal bovine serum (FBS, #10099141, Gibco), 1% (w/v) sodium pyruvate, 1% (w/v) penicillin/streptomycin, and 1% (w/v) non-essential amino acids (#11360070, #15140122, #11140050, Thermo-Fisher Scientific). Cells were kept for 10 days in 10 ng/mL GM-CSF to attain differentiation into naïve (unpolarized, CD68+) macrophages. Naïve macrophages were confirmed by flow cytometry using anti-CD68 antibodies (**Supplementary Figure S1C**). Naïve macrophages from each donor were split and the donor-paired fractions were used for polarization into either pro- or anti-inflammatory macrophages. Pro-inflammatory macrophages were obtained by incubating cells in FBS-free media supplemented with 20 ng/mL IFN-γ and 10 pg/mL LPS for 72 h. Anti-inflammatory macrophages were obtained by incubating cells in 20 ng/mL IL-4 and 20 ng/ml IL-13 for 72 h as described [50]. Robust polarization into pro-inflammatory (CD86+) and anti-inflammatory (CD206+) macrophages were confirmed by flow cytometry using anti-CD86 and anti-CD206 antibodies.

### Isolation of cell surface proteome, secretome and total cell lysate

Protein extracts from the 1) cell surface and cytosolic fractions, 2) secretome and 3) total cell lysates were isolated from the pro- and anti-inflammatory macrophage sub-populations.

1. Polarized macrophages (10^8^/sample) were washed with PBS and biotinylated by 1.2 mg EZ-Link Sulfo-NHS-SS-biotin for 45 min at room temperature using Pierce Cell Surface Protein Isolation kit (#A44390, Thermo-Fisher). Excess biotin reagent was removed by washing cells in chilled PBS. Cells were collected using a cell lifter (#3008, Corning) or after TrypLE express enzyme treatment (#12604021, Thermo-Fisher Scientific) and pelleted at 2,000 x *g* for 10 min. The pellet was resuspended in RIPA lysis buffer supplemented with complete protease inhibitor (#11836153001, Sigma Aldrich) and lysed at 4°C for 20 min with regular vortexing. Cell lysates were spun at 15,000 x *g* for 20 min, and the proteinaceous supernatant containing the biotinylated surface proteins were mixed with streptavidin-coated magnetic beads. After 90 min incubation at room temperature with end-over-end mixing, captured biotinylated (cell surface) proteins were isolated from the non-biotinylated (cytosolic) proteins by centrifuging the mixture through a spin column. Both biotinylated and non-biotinylated fractions were precipitated using acetone (1:4, v/v, lysate:acetone) at −30°C overnight. Tubes were centrifuged at 15,000 x *g* for 30 min, and precipitated pellets were washed with chilled acetone diluted three-fold in water. The protein extracts were pelleted at 15,000 x *g* for 15 min, dried and reconstituted in 8 M urea. Protein concentrations were determined by Qubit fluorometry (Thermo-Fisher Scientific).
2. The cytokine-containing polarization media was removed from fully polarized macrophages, and cells were washed and kept in serum-free RPMI media containing 1% pen-strep, 1% non-essential amino acid and 1% sodium pyruvate for 72 h. The media containing secreted proteins (secretome) was collected and left undisturbed for 30 min and centrifuged at 300 x *g* for 10 min to sediment cellular debris and particles. The secretome was concentrated using Amicon Ultra-10 kDa cutoff concentrator (Merck) and protein concentrations determined as above.
3. Polarized macrophages (10^8^/sample) were washed with cold PBS, collected using a cell lifter (#3008, Corning) or after TrypLE express enzyme treatment (#12604021, Thermo-Fisher Scientific) and pelleted at 800 x *g* for 15 min. Cells were lysed using RIPA buffer supplemented with complete protease inhibitor (#11836153001, Sigma Aldrich) in a shaker (1,000 rpm) for 20 min at 4°C, followed by centrifugation at 12,000 x *g* for 20 min at 4°C. The proteinaceous supernatant was transferred into fresh tubes and chilled neat acetone was added to precipitate protein overnight at −30°C. The protein extracts were pelleted, washed, dried, reconstituted in 8 M urea and the protein concentration determined as above.

### Glycomics

Macrophage protein extracts (lysate: 15 µg/sample; cell surface: 10 µg/sample) were reduced by 10 mM dithiothreitol (DTT) for 30 min, 37°C, and alkylated using 40 mM iodoacetamide for 30 min in dark. The reaction was quenched using excess DTT. The *N*-glycan release and desalting was performed as described [51]. Briefly, proteins were immobilized on a primed 0.45 µm polyvinylidene difluoride membrane (Merck-Millipore), stained with Direct blue 71 (Sigma-Aldrich), and excised and transferred to a flat bottom polypropylene 96-well plate (Corning). Membranes were blocked with 1% (w/v) polyvinylpyrrolidone in 50% (v/v) methanol and washed with water. *N*-glycans were released using 10 U peptide:*N*-glycosidase F (PNGase F) in 15 μL water, 16 h at 37°C, hydroxylated using 100 mM ammonium acetate, pH 5, 1 h at room temperature and reduced using 1 M sodium borohydride in 50 mM potassium hydroxide, 3 h at 50°C. Following *N*-glycan release, *O*-glycans were released from the same membranes using 0.5 M NaBH_4_ in 50 mM KOH for 16 h at 50°C. The reduction reaction was quenched using 2 μL glacial acetic acid. Dual desalting of the reduced *N*- and *O*-glycans was performed using strong cation exchange resin and then porous graphitized carbon (PGC) packed as micro-columns on top of C18 discs in P10 solid-phase extraction (SPE) formats. Glycans were eluted from the PGC SPE micro-columns using 0.1% trifluoroacetic acid (TFA) in 50% (both v/v) acetonitrile (ACN), dried, reconstituted in 10 μL water and transferred into high recovery glass vials (Waters) for LC-MS/MS analysis. Bovine fetuin was included as a reference to ensure efficient sample preparation and LC-MS/MS instrument performance.

Released glycan mixtures were characterized by PGC-LC-MS/MS. Samples were injected on a temperature-regulated (45-50°C) HyperCarb PGC LC capillary column (1 mm × 30 mm, 3 μm particle size, Thermo) installed on either an Agilent 1260 HPLC system or a Dionex UltiMate 3000 UHPLC system interfaced with either a LTQ Velos Pro linear ion trap (Thermo-Fisher Scientific) or an Amazon Speed ion trap (Bruker). For the Agilent 1260 HPLC – Thermo LTQ Velos Pro system, *N*-glycans were separated over a 60 min gradient of solvent A (10 mM ammonium bicarbonate) and solvent B (10 mM ammonium bicarbonate in 70 % (v/v) ACN): 0-3 min – 0 % B, 4 min – 14% B, 40 min – 40% B, 48 min – 56% B, 50-54 min – 100% B, 56–60 min – 0% B at a constant flow rate of 4 µL/min. The MS was operated in negative polarity mode with a HESI source (55°C), spray voltage 2.75 kV, sheath gas flow 13, auxiliary gas flow 7, capillary temperature 275°C. Full zoom MS1 scans were performed with a scan range of *m/z* 600-2,000, automatic gain control (AGC) of 3 x 10^4^ ions, three micro scans, and a maximum of 100 ms accumulation time. Top five precursor ions were selected for MS/MS (scan range *m/z* 200-2,000, AGC 1 x 10^4^ ions, maximum 100 ms accumulation time, isolation window *m/z* 1.4, and a normalized collision energy (NCE) of 33. For the Dionex UltiMate 3000 UHPLC – Bruker Amazon Speed ion trap setup, *N*-glycans were separated over a 65 min gradient: 0-5 min – 1 % B, 6 min – 14% B, 25 min – 25% B, 45 min – 70% B, 47-50 min – 98% B, 51–65 min – 1% B at a constant flow rate of 15 µL/min. The MS was operated with an ESI spray voltage 3.3 kV, nitrogen gas flow 6 L/min, dry temperature 207°C. MS1 scans were performed in the Ultra Scan mode with scan range of *m/z* 460-1,800, AGC of 7 x 10^4^ ions, 3 micro-scans and a maximum of 200 ms accumulation time. Top 3 precursor ions were selected for MS/MS (scan range *m/z* 100-2,200), AGC 3 x 10^4^ ions, isolation window *m/z* 4, and NCE 35. All MS and MS/MS data were acquired in profile mode and dynamic exclusion was disabled.

LC-MS/MS raw data were browsed using either Xcalibur v2.2 (Thermo-Fisher Scientific) or Compass Data Analysis v4.0 (Bruker). Glycans were manually identified based on monoisotopic molecular mass, MS/MS fragmentation pattern and PGC-LC retention time as described [52]. EIC based, area-under-the-curve glycan quantification was performed using Skyline v.23.1 [53, 54]. Relative *N*-glycan abundance was determined as a proportion of all *N*-glycans in each sample. Glycan features (e.g. α2,6-sialylation levels) were quantified by summing glycans carrying those structures. Glycans were depicted using GlycoWorkBench v2.1 according to the latest SNFG nomenclature [55].

### Proteomics

Macrophage protein extracts (cell surface: ∼2 μg/sample; cell lysate/secretome: 5 µg/sample) were reduced and alkylated as above. Sequencing grade porcine trypsin (#V5113, Promega) was added (3%, w/w, trypsin:protein) and mixtures were incubated overnight at 37°C. The reaction was stopped using 1% (v/v) TFA. The resulting peptide mixtures were desalted with hydrophilic-lipophilic-balanced SPE using home-made Oligo R3-C18 micro-columns and eluted sequentially with 50 μL 0.1% TFA in 50% (both v/v) ACN and then with 50 μL 0.1% TFA in 70% (both v/v) ACN. The eluted peptide fractions were combined, dried and reconstituted in 0.1% (v/v) formic acid for LC-MS/MS analysis.

Peptides were analyzed using a Q-Exactive HF-X Orbitrap operating in positive polarity mode and interfaced with an UltiMate 3000 nanoLC system (Thermo Scientific). Peptides (0.5 μg/sample) were loaded on a trap column (300 µm ID x 5 mm, Acclaim PepMap 100 C18 HPLC column, Thermo Fisher Scientific) and separated with a flow rate of 300 nL/min on an analytical nanoLC column (75 µm ID x 30 cm) packed in-house with ReproSil-Pur C18 column with AQ 3 µm resin (Dr Maisch, Germany). The mobile phases were 0.1% formic acid in 99.9% (both v/v) ACN (solvent B) and 0.1% (v/v) formic acid (solvent A). LC gradient: 2-33% B over 36 min, 33-95% B over 5 min, 95% B over 9 min, 95-2% B over 1 min, and kept at 2% B for 19 min. The Orbitrap acquired full MS1 scans with an AGC target of 3 x 10^6^ ions and a maximum fill time of 50 ms. MS1 scans were acquired at 60,000 resolution and a range of *m/z* 450-2,000. From each MS1 scan, the ten most abundant precursor ions (z ≥ 2) were selected for higher-energy collision-induced dissociation (HCD) fragmentation performed using NCE 28. Fragment ions were measured at 15,000 resolution with an AGC target of 2 x 10^5^ ions and maximum injection time of 28 ms using *m/z* 1.2 isolation windows and 30 s dynamic exclusion.

In addition to the data-dependent acquisition (DDA), specific ST6GAL1 peptides were measured using parallel reaction monitoring (PRM) using the same setup as above with minor modifications. The gradient increased from 2-30% B over 44 min, 30-95% B over 8 min, and 95-2% B over 8 min. Full MS1 scans were acquired at 120,000 resolution over an acquisition range *m/z* 350-1,200. HCD fragmentation was performed using a ST6GAL1-specific inclusion list with 30,000 resolution, and AGC target of 2 x 10^5^ ions and maximum injection time of 60 ms using an isolation window of *m/z* 1.2 and NCE 30.

HCD-MS/MS raw files (DDA) were imported into MaxQuant v2.2.0.0 to identify and quantify proteins. The Andromeda search engine was used to search data against the reviewed UniProtKB Human Protein Database (20,428 reviewed entries, September 2024) with a precursor and product ion mass tolerance of 20 ppm. Oxidation of methionine (15.99 Da) and protein N-terminal acetylation (42.01 Da) were selected as variable modifications. Carbamidomethylation of cysteine (57.02 Da) was set as a fixed modification. All identifications were filtered to <1% protein FDR. Proteins were quantified using label-free AUC-based quantitation employing the ‘match-between-run’ feature (0.7 min match time windows). Relative protein abundance was estimated based on iBAQ intensity values [56].

For quantitation of ST6GAL1 peptides, the PRM raw data were analyzed by MaxQuant v2.2.0.0 using ‘match-between-runs’ (0.7 min match time windows, 20 min alignment time windows). Relative ST6GAL1 abundance was determined based on normalized protein intensity (LFQ intensity). Statistical analysis was performed by Perseus v.2.0.7.0.

### Lectin flow cytometry

Polarized macrophages were washed and resuspended in FACS buffer containing PBS with 0.1% (w/v) BSA. Cells were incubated with fluorophore-conjugated lectins SNA (Cy3) and MAL-I (FITC) (20 μg/mL) for 40 min on ice or at 4°C. Controls were prepared by pre-incubating the lectins with or without 0.5 M α-lactose monohydrate before lectins were applied to the cells or by pre-treating cells with either broad (#P0722S, New England Biolabs) and α2,3-specific (#P0743S, New England Biolabs) neuraminidase for 45 min at 37°C. Following incubation with lectins, cells were washed with FACS buffer and filtered through a 100 µm FACS tube strainer. Lectin flow cytometry data was acquired on a CytoFLEX (Beckman Coulter) and data analyzed using the CytExpert software (Beckman Coulter).

### Lectin, biotin and actin-focused fluorescent microscopy

Polarized macrophages (∼3 x 10^4^/sample) were washed with cold PBS and fixed onto separate glass slides with chilled 4% (v/v) PFA, pH 7.4 for 15 min at room temperature. Slides were washed three times with cold lectin buffer (20 mM Tris HCL or base, 100 mM NaCl, 1 mM CaCl_2_, 1 mM MgCl_2_, and double distilled water, pH 7.4), incubated with 20 μg/mL SNA for 1 h at room temperature in the dark and then with 20 μg/mL MAL-I for 1 h at room temperature. Controls as above were included. Slides were briefly washed with TBS (1x) before 1 μg/mL DAPI was added and incubated for 10 min, then washed with TBS (1x). Antifade mounting media (Vectashield, Vector Laboratories) was applied before fitting the coverslip to preserve fluorescence. Images were taken using an Olympus IX83 widefield fluorescence microscope (Olympus DP80) with three channels: DAPI (ex: 345 nm; em: 445 nm), FITC (ex: 488 nm; em: 518 nm) and Cy3 (ex: 550 nm; em: 565 nm). Fiji software was used for quantification by applying threshold procedure to segment regions of interest. Images were deconvoluted using CellSens software to enhance signal clarity and reduce background.

Anti-inflammatory macrophages (∼3 x 10^4^/sample) were also incubated with 1.2 mg EZ-Link Sulfo-NHS-SS-biotin for 1 h followed by another 1 h incubation with streptavidin conjugated to Alexa Fluor™ 488 (#S32354, Thermo-Fisher Scientific). Images were captured using the same setup as above using the FITC channel. In separate experiments, ActinGreen488 ReadyProbe was used as per the manufacturer’s instructions (#R37110, Thermo-Fisher Scientific) to stain accessible actin. Three drops of ActinGreen488 ReadyProbe were added to pre-fixed macrophages (∼3 x 10^4^/sample), incubated for 45 min and washed with lectin buffer before adding SNA and DAPI as above. Images were captured using settings as above.

### Immuno- and lectin histochemistry

Formalin-fixed paraffin-embedded (FFPE) tissue sections (thickness: 5 µm/slide, size: 2 cm x 2 cm) of CRC patients were provided by Victoria Cancer Biobank (VCB). Ethics approval for the collection, handling, and analysis of the CRC tissue sections was obtained from Macquarie University HREC (5201800073). Tumor tissue sections paired with the corresponding normal adjacent tissues from unaffected parts of the colon from CRC patients with stage 4 disease were investigated using IHC following a published protocol with minor modifications [57]. Briefly, sectioned tissues on slides were incubated for 1 h at 50°C followed by three rounds of deparaffination in a neat xylene solution for 30 min at room temperature, rehydration over multiple steps in 70-100% (v/v) ethanol (3 min/step). Deparaffinated tissues were washed in TBS (1x) and antigens were heat-retrieved by incubating tissue slides in tris-ethylenetriaminetetracetic acid (EDTA) buffer containing 10 mM tris base, 1 mM EDTA, and 0.05% (w/v) Tween20, pH 9 for 30 min at 95°C. The antigen-retrieved tissue sections were cooled and washed three times (5 min each) with TBS (1x). The slides were blocked with 5% (w/v) BSA (Sigma, 99.9%, w/w, purity). BSA was pre-treated with 100 mM sodium periodate for 2 h at 4°C and dialyzed overnight in TBS (1x) to eliminate interfering glycoepitopes. Slides were washed with lectin buffer and incubated for 2 h after 20 μg SNA and MAL-I were added to different slides from the same tissue section. Slides were carefully rinsed with TBS (1x) for 3 min and incubated with anti-CD86 antibody (20 μg/mL) for 4 h, followed by anti-CD206 antibody (15 μg/mL) for another 4 h. DAPI (1 μg/mL) was added and slides incubated for 10 min, with gentle washing in TBS (1x) after each incubation step. Staining was performed at room temperature. The lectin and antibody staining were visualized, and co-localization assessed using an Olympus FV-3000 confocal microscope under 100x magnification using oil immersion objectives. Image data were collected and processed using CellSens imaging software (Olympus, Japan). Four different fluorescent channels were used to visualize staining: Ch1-DAPI 405 nm (detection wavelength; 430-470 nm), Ch2-Alexa flour 488 nm (500-540 nm), Ch3-Cy3 561 nm (570-620 nm), Ch4-Alexa flour 647 nm (650-750 nm). DAPI-cell nucleus, Alexa flour 488-CD206, Cy3-either SNA or MAL-I, Alexa flour 647-CD86. Co-localization of macrophage markers with lectins was quantified using the intensity-based ‘colocalization’ module within Imaris (v10.2.0, Oxford instrument). Intensity thresholds that best represent the signal levels for both channels were manually optimized. Pearson correlation co-efficient (PCC) was determined as follows to determine the degree of co-localization [58]:

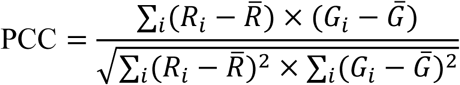

PCCs (r) above 0.70 were considered strong co-localization, 0.3-0.7 low-moderate co-localization and below 0.30 negligible co-localization [59].

### ST6GAL1 silencing

Accel siRNA (#A-010486-15-0020, Dharmacon) was used to target *ST6GAL1* for siRNA-based gene silencing using the open reading frame region. In brief, naïve macrophages (∼5 x 10^4^/sample) were pre-treated with and without 1.2 µM siRNA for 24 h, excess siRNA reagents removed, and cells then polarized into anti-inflammatory macrophages as per above. Cells were cultured for an additional 96 h before measuring ST6GAL1 protein levels in WT vs *ST6GAL1*-silenced conditions by western blotting. The target sequence for *ST6GAL1* silencing (5’ to 3’) was: GGUGGAUAUUUAUGAGUUC.

### Western blotting

Anti-inflammatory (WT and *ST6GAL1*-silenced) and pro-inflammatory macrophages (∼10^6^/sample) were washed in cold PBS and collected after TrypLE treatment. Cells were lysed using RIPA buffer and shaken for 10 min at 4°C. After centrifugation at 12,000 x *g* for 20 min at 4°C, proteins in the cell lysate fraction were quantified using Qubit and denatured with a SDS sample loading buffer (5x) for 5 min at 95°C. Proteins (50 µg/sample) were subjected to SDS-PAGE. After electrophoresis, proteins were transferred to a nitrocellulose membrane and blocked with 5% (w/w) oxidized BSA (see above). Mouse anti-ST6GAL1 (#H00006480-M01, Thermo) and mouse anti-β-actin (#A5316, Sigma) monoclonal antibodies were applied. A horseradish peroxidase-conjugated secondary anti-mouse antibody (#HAF007, R&D) was applied and signals detected with a chemiluminescent HRP substrate using ChemiDoc^TM^ XR (Bio-Rad). Images were acquired using ImageLab^TM^ software (Bio-Rad) and the relative protein levels quantified by ImageJ software (NIH).

### Live cell recording

Anti-inflammatory macrophages (WT and *ST6GAL1*-KOs) were cultured with and without LIM2405 cells (1:10, mol/mol, LIM2405:CD206+ macrophages). For both the mono- and co-cultures, the cells were kept for 72-96 h. During this period, live cell recordings were performed over a period of ∼21 h (mono-cultures) and ∼29 h (co-cultures) using an Olympus IX83 inverted microscope equipped with a zero-drift correction system and stage top incubator that maintained a constant temperature and CO_2_ level. One frame was captured every 5 min in a fixed field of view. Videos were generated using the CellSens Dimension software (Olympus). Cell motility was quantified by Fiji “manual tracking” plugin (Lopez et al., 2023) and cell interactions were manually counted.

### Sialyltransferase and SIGLEC gene expression

The expression of *ST6GALl* and select *SIGLEC* genes was assessed using cDNA micro-array chip datasets of M0 (naïve), M1 (pro-inflammatory macrophages), and M2 (anti-inflammatory macrophages) peripheral blood-derived macrophages available from NCBI Gene Expression Omnibus (GEO, GSE5099). Only M1 and M2 macrophage data were analyzed for *SIGLEC1*, *SIGLEC7* and *SIGLEC9* gene expression. Raw data were processed using GEO2R which normalized the gene expression level to log 2-fold change.

### Statistics

GraphPad prism v9.4.1 and Perseus v2.0.7.0 were used for statistical analyses. Volcano plots were made using Metaboanalyst v5.0. Significance was assessed using paired two-tailed student’s t-tests and non-parametric Mann-Whitney U tests or Wilcoxon tests, and Benjamini & Hochberg (false discovery rate) for pathway enrichment analysis. PCC raw data obtained from Imaris were interrogated using Microsoft Excel. *p* < 0.05 was used as significance threshold. Number of replicates, individual data points and confidence levels are indicated.

## 3. Results

### 3.1 Robust macrophage polarization

To gain insights into the dynamic glycosylation of polarizing macrophages, we firstly employed an *ex vivo* model system using peripheral blood-derived CD14+ monocytes isolated from buffy coats of four healthy donors to obtain naïve macrophages (**Supplementary Figure 1**). The naïve macrophages were polarized into pairs of pro- and anti-inflammatory macrophages (**Figure 1A**). Polarization of the macrophages was validated using microscopy (**Figure 1B-C**) and flow cytometry (**Figure 1D-E**), which confirmed the expected morphological and cellular characteristics of the two macrophage sub-populations. Consistent with their known morphological features [60], the relatively smaller pro-inflammatory macrophages (CD86+) exhibited lamellipodia-like structures whereas the comparably larger anti-inflammatory macrophages (CD206+) more often exhibited elongated filopodia. These data demonstrate that robust macrophage polarization was achieved.

**Figure 1.**
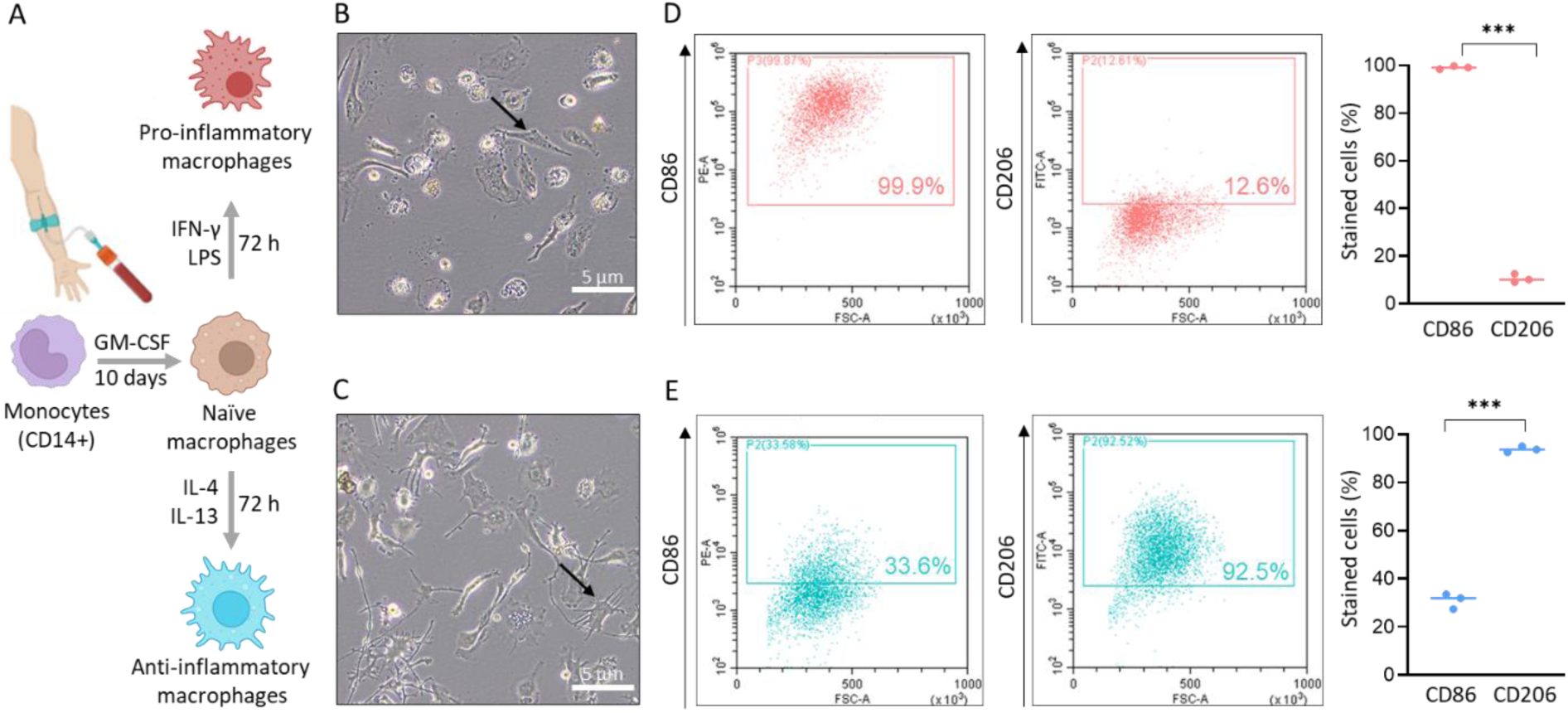
Robust polarization of monocyte-derived macrophages into pro- and anti-inflammatory macrophages. **A**) Monocytes were isolated from buffy coats of four healthy donors (n = 4). Monocytes were differentiated into naïve macrophages by GM-CSF (Supplementary Figure S1) and polarized into pairs of pro- and anti-inflammatory macrophage populations from each donor. For validation, bright field microscopy was used to validate the morphology of **B**) pro-inflammatory and **C**) anti-inflammatory macrophages captured with 10x magnification. Representative images are shown. Black arrows: Polarization-specific morphological traits are highlighted. Scale bars: 5 μm. Flow cytometry confirmed the expected expression patterns of known polarization-specific surface markers on the **D**) pro-inflammatory (CD86+ dominant) and **E**) anti-inflammatory (CD206+ dominant) macrophage populations. For D-E, two-tailed paired t-tests (\*\*\**p* < 0.001, n = 3).

### 3.2 Profound sialyl linkage switching in pro- and anti-inflammatory macrophages

Quantitative glycomics applied to lysates of the two macrophage sub-populations identified a total of 69 *N-*glycan structures across their cellular *N*-glycome (**Supplementary Table S1**). The pro- and anti-inflammatory macrophages exhibited similar *N*-glycan type distribution consisting mostly of complex-type *N*-glycans (40-45%) most of which were found to be sialylated structures (∼95%) (**Supplementary Figure S2A**). However, interrogating the PGC-LC-MS/MS glycomics data for glycosidic linkage features revealed that α2,6-sialylation was elevated and α2,3-sialylation reduced in the anti-inflammatory relative to pro-inflammatory macrophages (*p* < 0.01) (**Supplementary Figure S2B**). Consistent sialyl linkage differences were observed for the two macrophage sub-populations across all four donors as exemplified by a prominent biantennary disialylated *N*-glycan (*m/z* 1184.4) spanning three different sialo-isomers (**Supplementary Figure S2C**). For these abundant *N*-glycan isomers, the α2,6/α2,3-sialyl linkage ratio was significantly higher for the anti-inflammatory (30.5 ± 14.6) relative to the pro-inflammatory (7.1 ± 6.9, *p* = 0.0155) macrophages.

As sialylation is a common cell surface feature known to impact immune cell function and interactions [61–64], we investigated the sialyl linkage distribution on the surface of the two macrophage sub-populations using an established biotinylation approach [65]. Proteomics was used to validate the capture of known cell surface proteins such as HLA, integrins, ITGAM, and MRC1 in the biotinylated fractions (**Supplementary Table S2-3**). Pathway enrichment analysis indeed confirmed a significant enrichment of cell surface proteins in the biotinylated fractions (e.g. plasma membrane FDR = 6.81 x 10^-47^) relative to non-biotinylated fractions, which instead mostly contained cytosolic proteins (e.g. cytosol FDR = 9.56 x 10^-64^) (**Supplementary Figure S3**).

Focusing again on the biantennary disialylated *N*-glycan isomers (*m/z* 1184.4), a dramatic difference in the sialyl linkage distribution was observed on the cell surface with the α2,6/α2,3-sialyl linkage ratio consistently higher for the anti-inflammatory (57.3 ± 21.4) relative to pro-inflammatory (3.9 ± 0.9, *p* = 0.0153) macrophages across all donors (**Figure 2A**). Expanding the analysis to encompass all major surface sialo-*N*-glycans (NeuAc_1-2_Gal_2_Man_3_GlcNAc_4_Fuc_0-1_), a profound sialyl linkage switching was observed with raised α2,6-sialylation (87.6% ± 1.9%) and severely depleted α2,3-sialylation (1.2% ± 0.4%) being key features of the surface *N*-glycans of anti-inflammatory macrophages relative to those of pro-inflammatory macrophages that expressed comparably less α2,6-sialylation (50.3% ± 5.1%) and more α2,3-sialylation (10.1% ± 0.9%) (both *p* < 0.001) (**Figure 2B** and **Supplementary Table S4**).

**Figure 2.**
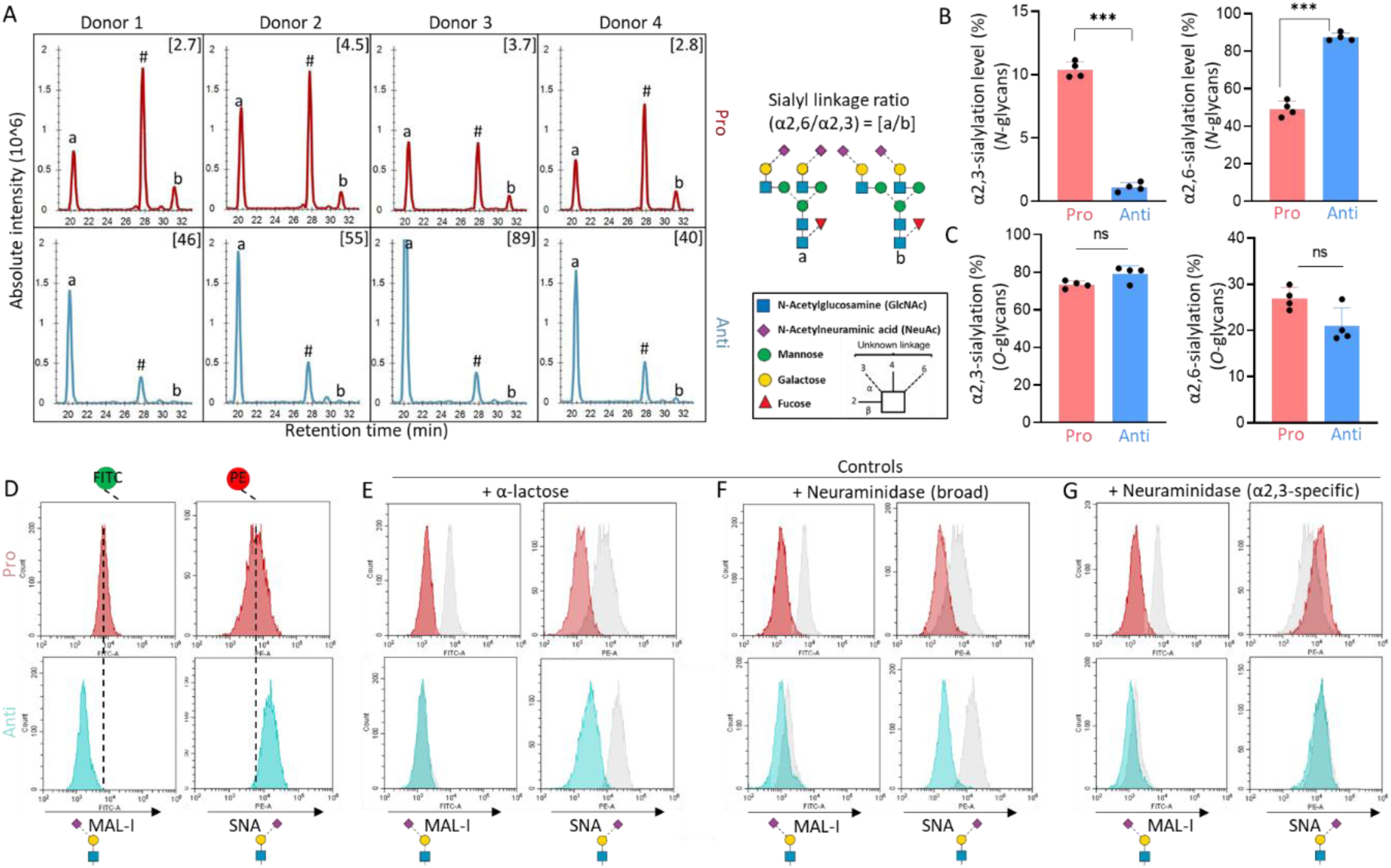
Profound sialyl linkage switching on the cell surface of pro- and anti- inflammatory macrophages. **A**) Extracted ion chromatograms of PGC-LC/MS/MS *N*-glycomics data of the macrophage cell surface fraction focusing on a prominent biantennary disialylated *N*-glycan (*m/z* 1184.4) (Hex_5_HexNAc_4_dHex_1_NeuAc_2_) spanning three discrete sialo-isomers for pro-inflammatory (red, upper row) and anti-inflammatory (blue, lower row) macrophages. The sialo-isomers carried both α2,6-linked (a), α2,3-linked (b) and a combination of both α2,6/α2,3-linked (marked #) sialylation, only a-b isomers were used to determine the α2,6/α2,3-sialyl linkage ratio. Glycans are depicted according to SNFG nomenclature [55]. **B**) Quantitation of α2,3- (left) and α2,6- (right) sialylation levels based on PGC-LC/MS/MS *N*-glycomics data of abundant *N*-glycans from the macrophage cell surface fractions. **C**) Quantitation of α2,3- (left) and α2,6- (right) sialylation levels based on PGC- LC/MS/MS *O*-glycomics data of abundant *O*-glycans from the macrophage cell surface fractions (see also Supplementary Figure S4A-B). See Supplementary Table S4-S5 for surface *N*- and *O*-glycome data. For B-C, two-tailed paired t-tests (\*\*\**p* < 0.001, ns: non-significant, n = 4). For both graphs, data are presented as mean ± SD, individual data points are shown. **D**) MAL-I (α2,3-sialic acid reactive) and SNA (α2,6-sialic acid reactive) based lectin flow cytometry of pro-inflammatory (upper row, red) and anti-inflammatory (lower row, blue) macrophages. Controls were carried out including **E**) pre-treatment of lectins with α-lactose, and pre-treatment of investigated cell populations with **F**) neuraminidase (broad) and **G**) α2,3-specific neuraminidase. See Supplementary Figure S5 for quantitation of SNA and MAL-I reactivity across all conditions.

We then explored whether sialyl linkage switching was also mirrored in the surface *O*-glycome of the macrophage sub-populations. Global PGC-LC-MS/MS-based linkage profiling revealed no significant differences in the sialyl linkage distribution in the surface *O*-glycans of pro- and anti-inflammatory macrophages (**Figure 2C** and **Supplementary Table S5**). Some of the individual surface *O*-glycans even displayed a modest opposite switch in the sialyl linkage distribution as observed for a trisaccharide sialo-*O*-glycan exhibiting a mildly higher α2,6/α2,3-sialyl linkage ratio for the pro-inflammatory relative to anti-inflammatory macrophages (*p* < 0.01) (**Supplementary Figure S4A-B**). Taken together, the quantitative glycomics data demonstrate a profound sialyl linkage switching on the *N*-glycans (rather than the *O*-glycans) within the glycocalyx of the polarized macrophage sub-populations.

To validate the surface glycomics data, we used a sialic acid-focused lectin flow cytometry approach employing *Sambucus nigra* agglutinin (SNA, preferentially recognizing α2,6-sialylation) and *Maackia amurensis* lectin I (MAL-I, preferentially recognizing α2,3-sialo-epitopes) [66]. Consistent with the *N*-glycomics data, MAL-I stained predominantly the pro-inflammatory macrophages indicating a severe depletion of α2,3-sialylation on the surface of anti-inflammatory macrophages (**Figure 2D**, see **Supplementary Figure S5A** for quantitation). Conversely, the anti-inflammatory macrophages were predominantly recognized by SNA. Key controls were carried out to confirm the lectin reactivity. Firstly, we used α-lactose, a disaccharide known to competitively inhibit both SNA and MAL-I [67]. The pre-treatment of lectins with α-lactose led to a noticeable shift back to the baseline in all conditions except for the already insignificant MAL-I staining of anti-inflammatory macrophages thereby confirming carbohydrate-specific lectin recognition (**Figure 2E** and **Supplementary Figure S5B**). Similarly, pre-treatment of cells with broad neuraminidase reduced the lectin staining in both macrophage sub-populations confirming sialic acid-mediated recognition by both SNA and MAL-I (**Figure 2F** and **Supplementary Figure S5C**). Importantly, pre-treatment of cells with α2,3-specific neuraminidase exclusively decreased the MAL-I recognition of pro-inflammatory macrophages validating the low α2,3-sialylation levels on the surface of anti-inflammatory macrophages (**Figure 2G** and **Supplementary Figure S5D**). As expected, the α2,3-specific neuraminidase did not affect the SNA staining of the anti-inflammatory macrophages and in fact mildly increased the SNA recognition of pro-inflammatory macrophages presumably because of better accessible to the available α2,6-sialyl glycoepitopes in the absence of α2,3-sialylation. Collectively, the surface glycomics and lectin flow cytometry data provide strong evidence of a dramatic sialyl linkage switching within the surface *N*-glycome of pro- and anti-inflammatory macrophages.

### 3.3 Proteinaceous nanotubes rich in α2,6-sialylation protrude from and form homotypic connections between anti-inflammatory macrophages

Next, we set out to visualize the spatial distribution of the sialo-glycoepitopes on the macrophage subtypes using SNA- and MAL-I-based lectin fluorescence microscopy. Consistent with the previous observations, the pro-inflammatory macrophages showed recognition by both SNA and MAL-I (**Figure 3A**). In this cell population, SNA stained the long filopodia-like structures that facilitate cell migration while MAL-I stained other regions of the macrophage cell surface including the lamellipodia as supported by limited co-localization of the two lectins. Expectedly, the anti-inflammatory macrophages were devoid of MAL-I recognition, but, surprisingly, exhibited extended and highly branched (yet well-coordinated) nanotubes rich in α2,6-sialylation as evaluated by their intense SNA reactivity (**Figure 3B**). The SNA-reactive branched nanotubes were sensitive to broad neuraminidase, but not to α2,3-specific neuraminidase. The nanotubes appeared as elongated branched protrusions from the plasma membrane often forming homotypic connections between adjacent anti-inflammatory macrophages. Consistent with the surface glycomics and lectin flow cytometry data, quantitation of the cell areas stained by lectin fluorescence confirmed that the anti-inflammatory macrophages were higher in α2,6-sialylation and severely depleted in α2,3-sialylation relative to the pro-inflammatory macrophages (both *p* < 0.001) (**Supplementary Figure S6A**). Upon broad neuraminidase treatment, the regions stained by SNA and MAL-I were significantly reduced in size in both macrophage sub-populations whereas only MAL-I staining was reduced upon α2,3-specific neuraminidase confirming the expected lectin recognition specificity (**Supplementary Figure S6B-C**).

**Figure 3.**
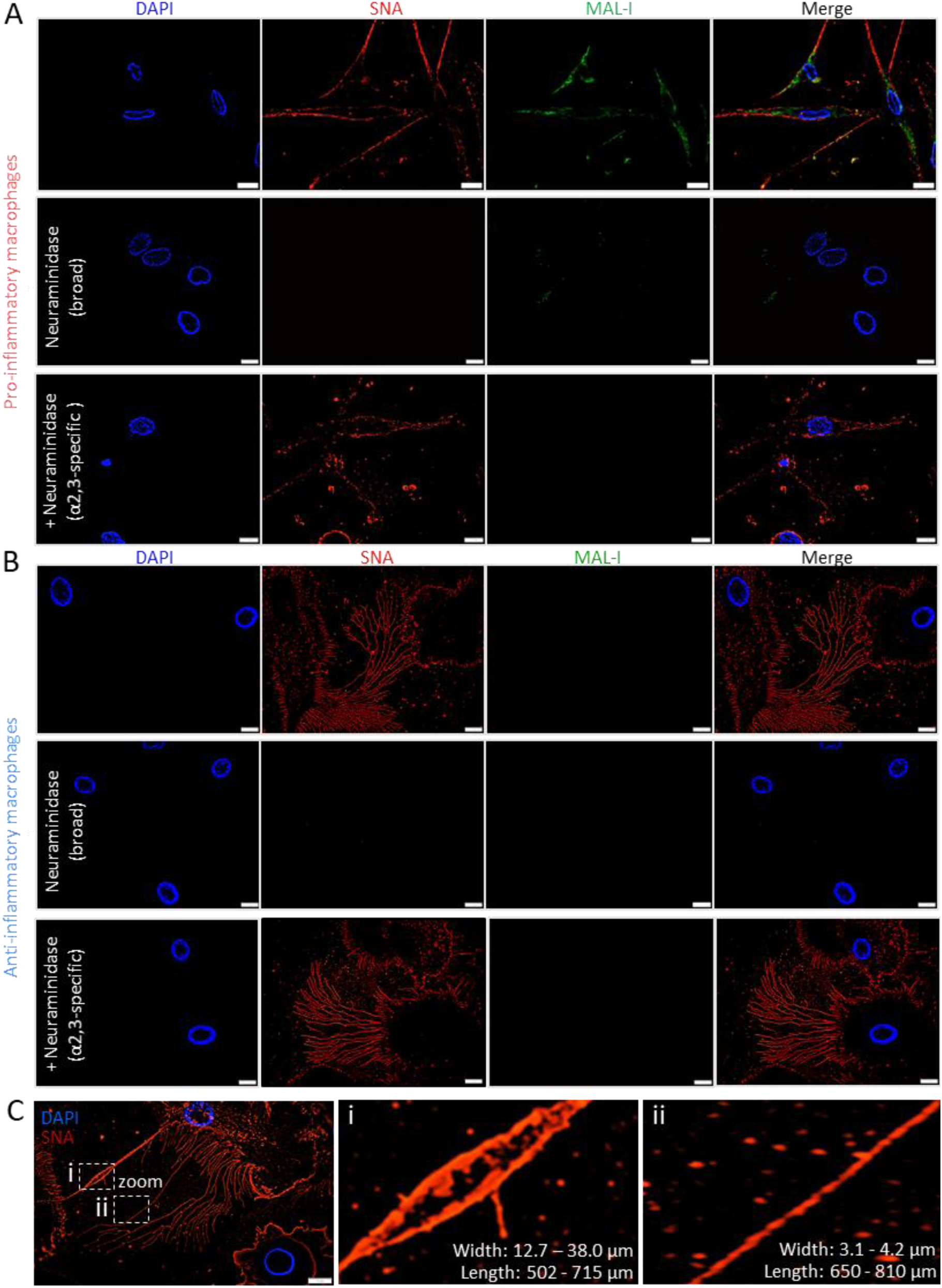
Branched α2,6-sialylated nanotubes form extensive networks that protrude from anti-inflammatory macrophages. SNA (α2,6-sialic acid reactive, red) and MAL-I (α2,3-sialic acid reactive, green) based lectin fluorescence microscopy of **A**) pro-inflammatory and **B**) anti-inflammatory macrophages. Nuclei were counterstained by DAPI (blue). Merged images were generated by combining all three channels. Representative images are shown from biological triplicates (n = 3). Pre-treatments of the investigated cell populations were performed with broad neuraminidase and α2,3-specific neuraminidase to ensure lectin recognition specificity. Scale bars: 10 µm. See Supplementary Figure S6A-C for quantitation of SNA and MAL-I reactivity across conditions. **C**) SNA-based lectin fluorescence microscopy of different types of α2,6-sialylated nanotubes in anti-inflammatory macrophages with zoomed regions focusing on (i) thick linear nanotubes (width: 12.7 – 38.0 μm, length: 502 – 715 μm) and (ii) thin branched (width: 3.1 – 4.2 μm, length: 650 – 810 μm). Length and width were measured by Fiji software.

As there, to the best of our knowledge, are only scattered reports of cellular nanotubes [68–71], we investigated the characteristics of the intriguing α2,6-sialylated nanotubes protruding from the anti-inflammatory macrophages. Two distinct types of nanotubes were observed: a group of thick and rather linear nanotubes (width: 12.7–38.0 µm) and another more prevalent group of thin and highly branched nanotubes (width: 3.1–4.2 µm) (**Figure 3C**). Although highly heterogeneous and dynamic, the two types of nanotubes exhibited similar length characteristics reaching up to 700-800 µm and both types were dynamically contracting and expanding over time. The thin and branched nanotubes appeared to serve as structural scaffolds for the thicker linear nanotubes to facilitate connections between adjacent cells. While the thicker nanotubes were observed for most anti-inflammatory macrophages, thin nanotubes appeared restricted to motile cells (see live cell recordings below).

### 3.4 ST6GAL1 drives cancer-promoting nanotubes carrying α2,6-sialylation within the anti-inflammatory macrophage population

Seeking to firstly identify glyco-enzyme(s) responsible for the sialyl linkage switching we used quantitative proteomics to profile the expression of sialyltransferases (STs) across the pro- and anti-inflammatory macrophages. Remarkably, ST6GAL1 that is known to catalyze α2,6-sialylation of *N*-glycans rather than *O*-glycans [72–74] was exclusively and consistently detected in the lysates of anti-inflammatory and absent (or present below quantitation levels) in the lysates of pro-inflammatory macrophages (*p* < 0.05) (**Figure 4A**, see also **Supplementary Table S6** for data). In support, ST6GAL1-focused proteomics of the macrophage secretome (**Figure 4B**) and western blot analysis of macrophage lysates (**Figure 4C**) also showed substantial elevation of ST6GAL1 in anti-inflammatory compared to pro-inflammatory macrophages. Finally, re-interrogation of public transcriptomic data from various macrophage sub-populations confirmed a higher *ST6GAL1* gene expression in anti-inflammatory compared to pro-inflammatory macrophages (*p* < 0.001) (**Figure 4D**). Other STs were identified in the transcriptomics data, but these were either relatively lowly expressed (*ST3GAL2*, *ST3GAL3*), or not relevant to *N*-glycans (*ST3GAL5*, *ST3GAL6*). Collectively, the data suggest that ST6GAL1 is strongly regulated upon macrophage polarization and therefore may contribute to the sialyl linkage switching of the surface *N*-glycome in pro- and anti-inflammatory macrophages.

**Figure 4.**
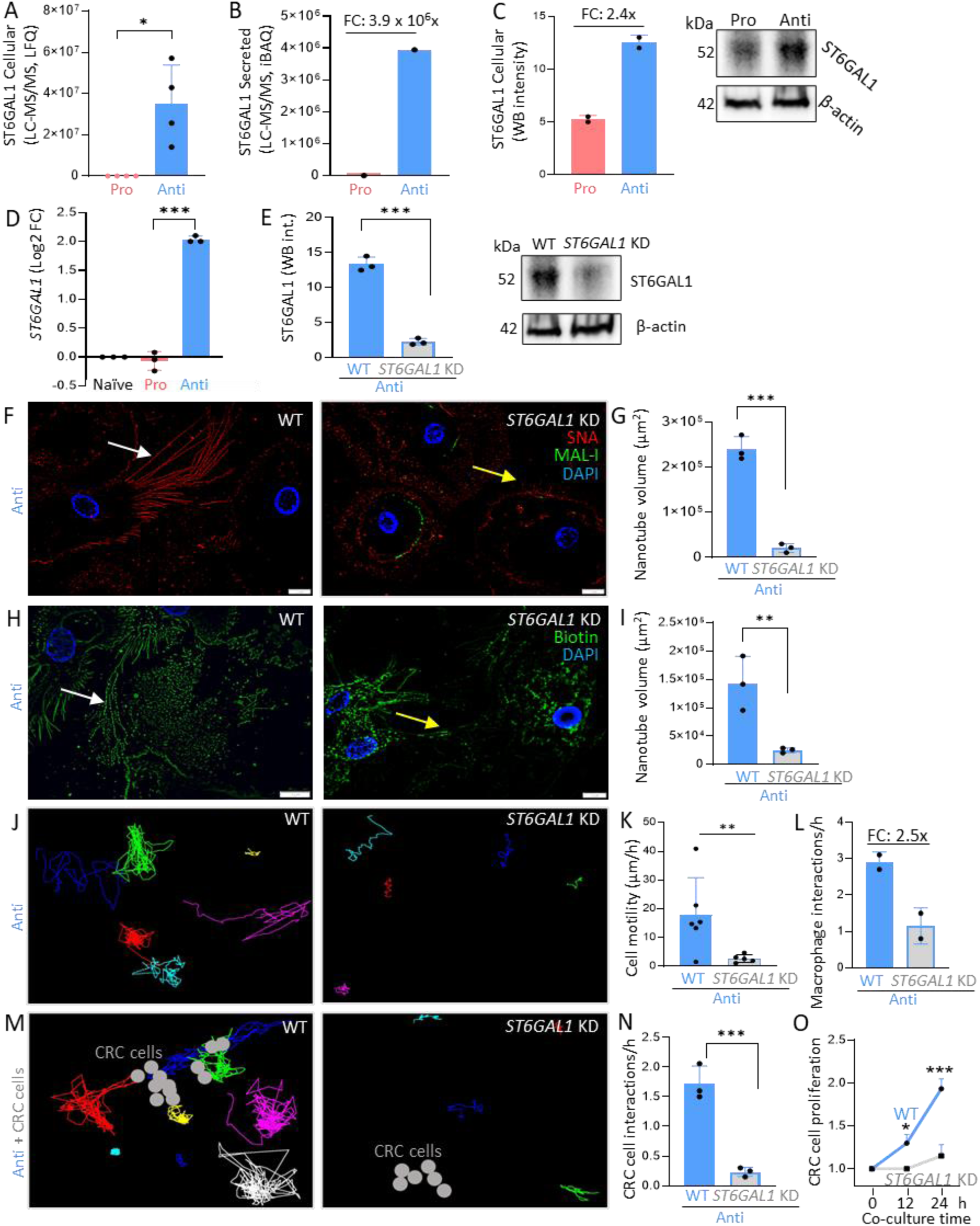
ST6GAL1 drives sialyl linkage switching and reduce nanotube protrusions, interactions and motility of anti-inflammatory macrophages. ST6GAL1 levels measured in the **A)** cellular (lysate) and **B**) secretome of pro- and anti-inflammatory macrophages using parallel reaction monitoring (PRM) of six ST6GAL1 peptides (n = 4) and untargeted proteomics (n = 1), respectively. FC: fold change. **C**) Western blot (WB) based quantitation of ST6GAL1 levels in lysates of pro- and anti-inflammatory macrophages by densitometry. Levels normalized to β-actin. FC: Average fold change (n = 2). **D**) *ST6GAL1* gene expression based on public transcriptomic data (GEO: GSE5099). Differences in transcript levels are presented as Log2 fold change (n = 3). **E**) WB-based quantitation of ST6GAL1 in untreated (WT) and *ST6GAL1*-knockdown (KD) anti-inflammatory macrophages based on densitometry (n = 3). Levels normalized to β-actin. **F**) SNA- (red) and MAL-I- (green) based lectin fluorescence microscopy of WT and *ST6GAL1*-KD anti-inflammatory macrophages. Representative images from biological triplicates (n = 3). DAPI in blue. **G**) SNA-based quantitation of the nanotube volume (µm^2^). **H**) Biotin fluorescence microscopy of non- permeabilized WT and *ST6GAL1*-KD anti-inflammatory macrophages visualized by streptavidin-FITC (green). Nuclei were counterstained by DAPI (blue). For F and H, scale bars are 10 µm and white and yellow arrows provide examples of intact and fragmented nanotubes, respectively. **I**) Quantitation of biotin-reactive nanotube volume (µm^2^) from three different images. Live cell recordings (21 h) of WT and *ST6GAL1*-KD anti-inflammatory macrophages in monocultures tracking **J**) the migration pattern of individuals cells (in different colors) (see also Supplementary Video S1) and **K**) motility measured as distance travelled by each cell over time (µm/h) in a single representative field (n = 5-6) out of two surveyed regions and **L**) homotypic cell interactions as manually counted and averaged over the live cell recording from biological duplicates. Live cell recordings (29 h) of WT and *ST6GAL1*-KD anti-inflammatory macrophages co-cultured with CRC cells (LIM2405, grey dots) tracking **M**) the migration pattern of individuals cells (in different colors) (see also Supplementary Video S2), **N**) heterotypic interactions with CRC cells and **O**) relative CRC cell proliferation. For N-O, data were obtained from three independent live cell recording experiments (n = 3). For all graphs, data are presented as mean ± SD, individual data points are shown and number of replicates indicated. Statistical tests were performed using paired/unpaired two-tailed t-tests, parametric tests, and non-parametric Mann-Whitney tests (\*\*\**p* < 0.001, \*\**p* < 0.01, \**p* < 0.05, ns, not significant *p* ≥ 0.05).

To confirm the biosynthetic link between ST6GAL1 and sialyl linkage switching and to explore any functional links between ST6GAL1, the α2,6-sialylated nanotubes and other phenotypic characteristics of anti-inflammatory macrophages, we performed siRNA-based *ST6GAL1* knockdown (KD) within this cell population. Robust *ST6GAL1* silencing was achieved as demonstrated by a significant protein depletion in *ST6GAL1* KO cells relative to ST6GAL1 levels in untreated (WT) anti-inflammatory macrophages (*p* < 0.001) (**Figure 4E**). Interestingly, the nanotubes appeared structurally disintegrated upon ST6GAL1 disruption as visualized by SNA lectin microscopy (**Figure 4F**). The intense SNA reactivity observed continuously along the protruding nanotubular structures in untreated anti-inflammatory macrophages was noticeable fragmented and reduced in intensity upon ST6GAL1 disruption resulting in a measurable decrease in nanotube volume (*p* < 0.001) (**Figure 4G**). While absent in untreated cells, MAL-I staining was visible in *ST6GAL1* KD cells presumably due to the loss of α2,6-sialylation previously masking underlying α2,3-sialo-epitopes possibly carried by shorter (less accessible) *O*-glycans in the glycocalyx (Figure 2C).

Using biotin fluorescence microscopy without prior membrane permeabilization to broadly stain accessible (surface) proteins, biotin-reactive proteins spanning the length of the nanotubular structures were observed in untreated (WT) anti-inflammatory macrophages with only minor distal regions of the nanotubes not staining continuously for biotin (**Figure 4H**). Similar to the SNA fluorescence microscopy, biotin staining indicated truncated and structurally fragmented nanotubes localizing around rather than protruding away from the cell body upon ST6GAL1 disruption resulting in a measurable reduction in nanotube volume compared to untreated anti-inflammatory macrophages (*p* < 0.01) **(Figure 4I**).

Given that the actin cytoskeleton plays a crucial role in cell migration and protrusion formation [75, 76], we tested if actin is a structural component of the observed nanotubes. However, actin-focused fluorescence microscopy did not provide evidence of actin within the nanotubular structures instead showing actin localizing in and around the cell body of anti-inflammatory macrophages (**Supplementary Figure S7**). Our data therefore suggest that actin-free nanotubes in anti-inflammatory macrophages carry exposed *N*-glycoproteins displaying α2,6-rather than α2,3-sialylation and indicate that ST6GAL1 is involved in the formation and/or stabilization of the α2,6-sialylated nanotubes.

Macrophage migration and communication are essential components of an effective innate immune response [77]. We therefore evaluated the impact of ST6GAL1 disruption on motility and cell interactions in anti-inflammatory macrophages. Macrophage migration was tracked using live cell recordings of anti-inflammatory macrophages in monocultures over 21 h (**Figure 4J** and **Supplementary Video S1**). Cell tracking revealed distinct motility differences with untreated (WT) anti-inflammatory macrophages averaging ∼20 µm/h, while ST6GAL1 disrupted cells exhibited severely impaired motility averaging <5 µm/h (*p* < 0.01) (**Figure 4K**). The impaired motility and nanotube disintegration caused by the ST6GAL1 disruption were accompanied by distinctly fewer homotypic interactions between the anti-inflammatory macrophages (∼1 interaction/h) relative to untreated cells (∼2.5-3 interactions/h) (**Figure 4L**).

Anti-inflammatory macrophages are intimately involved in supporting cancer cell growth and metastasis in the tumor microenvironment (TME) [78], a phenomenon particularly well reported in colorectal cancer (CRC) [23]. We therefore carried out co-cultures of anti-inflammatory macrophages and CRC cells (LIM2405) to examine the impact of ST6GAL1 disruption on macrophage motility and their heterotypic interactions to CRC cells as well as the proliferation rate of CRC cells using live cell recordings over a 29-h period (**Supplementary Video S2**). Similar to observations from the macrophage monocultures, the migration and motility of the anti-inflammatory macrophages were significantly impaired upon ST6GAL1 disruption in the presence of CRC cells (*p* < 0.01, quantitative data not shown) (**Figure 4M**). In addition, ST6GAL1 disrupted cells displayed consistently fewer heterotypic interactions with the CRC cells (∼0.2 interactions/h) compared to untreated anti-inflammatory macrophages (∼1.7 interactions/h, *p* < 0.001) (**Figure 4N**). Unlike the anti-inflammatory macrophages, the severely truncated nanotubes in the *ST6GAL1*-silenced cells were not able to form heterotypic connections to the adjacent CRC cells (**Supplementary Figure S8** and **Supplementary Video S3**). Possibly due to their reduced interactions with ST6GAL1-disrupted macrophages, the CRC cells in those conditions appeared to be less adhesive (more prone to detach from substrates) and were more easily phagocytosed than CRC cells co-cultured with untreated anti-inflammatory macrophages. Consequently, the CRC cells exhibited a considerably reduced proliferation rate when co-cultured with ST6GAL1-disrupted anti-inflammatory macrophages (**Figure 4O**) (estimated doubling time: ∼110 h) compared to the proliferation rate of untreated cells (doubling time: ∼24 h) (*p* < 0.001) (**Supplementary Figure S9**). This observation suggests a critical role of ST6GAL1 in anti-inflammatory macrophages in terms of supporting CRC cell adhesion and proliferation through sialic acid-rich nanotubule-mediated interactions.

### 3.5 Strong Siglec remodeling in polarizing macrophages

To probe how ST6GAL1 and α2,6-sialylation mechanistically alter macrophage interactions and communication, we performed quantitative proteomics of the cell lysate and surface fractions of pro- and anti-inflammatory macrophages. In total, 189 and 355 proteins were found to differ in abundance in the cell lysate and surface fractions, respectively, across the two macrophage sub-populations (*p* < 0.05) (**Supplementary Table S6**). Confirming the robustness of the proteomics data, MRC1, a known protein receptor of anti-inflammatory macrophages [79], was found to be elevated in that sub-population (FC: 10.9x, *p* = 1.6 x 10^-5^), while CD86, a marker of pro-inflammatory macrophages [80] indeed was elevated in the pro-inflammatory cell population (FC: 11.4x, *p* = 5.8 x 10^-3^) (**Supplementary Figure S10A-B**).

In addition to ST6GAL1 (see 3.4), amongst the most differentially expressed proteins were several sialic acid-binding immunoglobulin-like lectins (Siglecs), a large family of immune cell-derived signaling receptors known to recognize bespoke sialoglycans (Siglec ligands) on cell membranes [81]. Notably, Siglec-7 was found to be consistently elevated in the anti-inflammatory macrophages whereas Siglec-1 (lysate) and Siglec-10 (surface) were elevated in pro-inflammatory macrophages (**Supplementary Figure S11**). Transcriptomics confirmed the increased gene expression of *SIGLEC1* in pro-inflammatory and *SIGLEC7* in anti-inflammatory macrophages (all *p* < 0.01). Furthermore, proteome analysis of the secretome identified Siglec-7 exclusively in anti-inflammatory macrophages supporting that Siglec-7 may be important for those cell populations. These observations associate Siglec-7, Siglec-1 and Siglec-9 with the ST6GAL1-mediated (and thus α2,6-sialylation-driven) cellular communication and other glycophenotypic traits in anti-inflammatory macrophages.

### 3.6 Sialyl linkage switching of pro- and anti-inflammatory TAMs in CRC tumor tissues

Sialylation plays a critical role in progression, immune evasion and metastasis in CRC [33, 61, 82]. While many aspects remain unknown, the regulatory role of sialic acids are, at least in part, related to their ability to modulate (via Siglecs) the interactions between CRC cells and the infiltrating immune cells including the tumor-associated macrophages (TAMs) that are prevalent in CRC tumor tissues [83]. We therefore sought to investigate if the sialyl linkage switching observed in pro- and anti-inflammatory macrophages was reflected in TAMs residing in CRC tumor tissues. For this purpose, we used MAL-I and SNA-based lectin histochemistry and macrophage-focused IHC on CRC tissues resected from patients suffering from advanced CRC (stage 4). Consistent with our previous findings, MAL-I (α2,3-sialyl reactive) showed strong co-localization with pro-inflammatory TAMs (CD86+) (r > 0.7) and weak co-localization with anti-inflammatory TAMs (CD206+) (r < 0.1, *p* < 0.001) (**Figure 5A**). In contrast, SNA (α2,6-sialyl reactive) showed strong co-localization with anti-inflammatory TAMs (r > 0.7) and less with the pro-inflammatory TAMs (r < 0.3, *p* < 0.01) (**Figure 5B**). To validate these findings and control for any background staining, we surveyed adjacent normal colonic tissue from the same CRC patients. As expected, normal colonic tissues showed negligible levels of the macrophage markers and only poor MAL-I or SNA reactivity, none of which appeared to co-localize (**Supplementary Figure S12**). These observations document an absence of TAMs in the normal adjacent colonic tissue, which exhibited markedly less sialylation relative to the prominent MAL-I and SNA reactivity of the CRC tumor tissues. Taken together, these data indicate that the profound sialyl linkage switching observed in pro- and anti-inflammatory macrophages is recapitulated in TAMs residing within the CRC TME.

**Figure 5.**
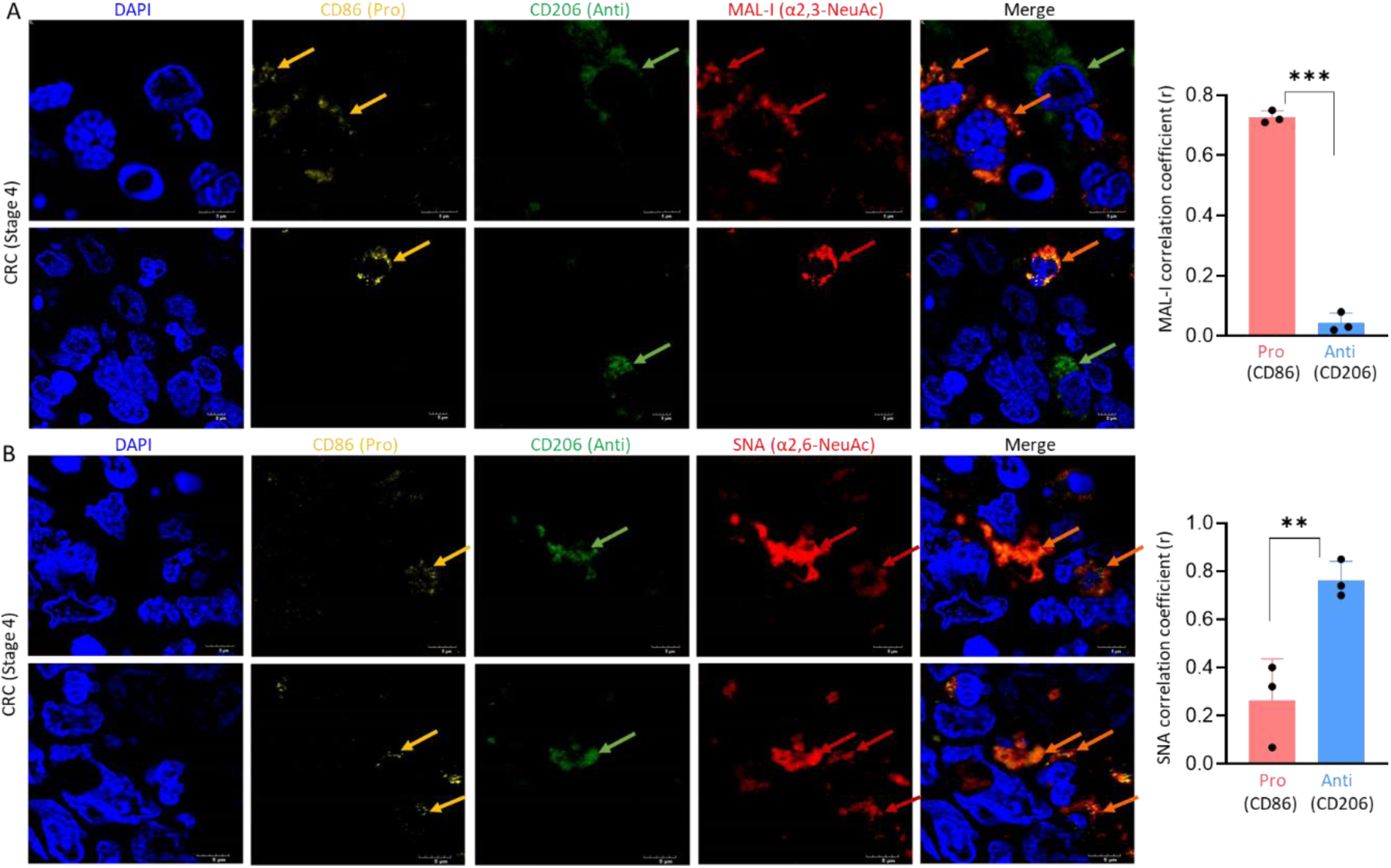
Sialyl linkage switching in TAMs residing in CRC tumor tissues. **A)** SNA- and **B**) MAL-I-based lectin histochemistry and immunohistochemistry (IHC) of tumor tissues resected from patients with advanced CRC (stage 4). Tissue sections were deparaffinized and stained with DAPI (nuclei), markers for pro-inflammatory (CD86, yellow) and anti-inflammatory (CD206, green) macrophages and lectins SNA (α2,6-sialic acid reactive) and MAL-I (α2,3-sialic acid reactive). Merged images were generated by combining all four channels. Arrows: Examples of unique or co-localizing features. Scale bars: 5 μm. For A-B, two representative images have been shown from three different tissue sample slides (n = 3). Right: Pearson correlation coefficients (r) of the polarization markers (CD86 and CD206) with MAL-I (A) and SNA (B). Two-tailed paired t-tests (\*\*\**p* < 0.001, \*\**p* < 0.01). For all graphs, data are presented as mean ± SD, individual data points are shown.

## 4. Discussion

Guided by chemical cues in the local tissue environment, macrophages polarize into pro- or anti-inflammatory phenotypes that play different functional roles in chronic inflammation, infection and cancer [19]. During the progression of colorectal cancer (CRC), for example, the tumor-fighting pro-inflammatory tumor-associated macrophages (TAMs) dominate the early disease stages, while tumor-supporting anti-inflammatory TAMs gradually become prevalent with advanced CRC and associate with poor patient outcome [23]. Despite an intense focus and recognized functional importance [84–86], the glycobiology underpinning the cellular plasticity and effector functions of TAMs remain incompletely described, leaving an important knowledge gap that prevents us from effectively manipulating macrophages from disease promoting towards disease fighting phenotypes and making use of their plasticity and potent biological effector functions as a therapeutic avenue.

Protein glycosylation is increasingly recognized to play important roles across a broad range of immunological processes associated with cancer [87, 88]. Unsurprisingly, there is therefore growing interest in elucidating the macrophage glycobiology to understand how glycans dynamically shape macrophage effector functions within the tumor microenvironment (TME). While macrophage glycosylation is known to influence cell signaling, adhesion and interactions with surrounding cells [89] and drive immune evasion [48], the glycocalyx remodeling accompanying TAM polarization remains poorly defined. Aiming to detail how glycans impact TAM effector functions, in this study we set out to define with high structural and spatial resolution the distinct glycophenotypes of pro- and anti-inflammatory macrophages.

Enabled by cell surface-focused glycomics, proteomics and lectin flow cytometry applied to homogenous and donor-paired populations of pro- and anti-inflammatory macrophages, our study revealed profound sialyl linkage switching within the surface *N*-glycome upon macrophage polarization. The surface *N*-glycans of the anti-inflammatory macrophages overwhelmingly displayed α2,6-sialylation and were near-absent in α2,3-sialylation, characteristics that were strikingly different from the surface *N*-glycome of pro-inflammatory macrophages carrying considerable levels of α2,3-sialylation. The rich biochemical details afforded by the applied -omics methods paired with spatial approaches and key controls (exoglycosidase enzymes and molecular competitors) for sialic acid linkage recognition, offered deep and quantitative insight into polarization-induced modulations of the macrophage glycocalyx and robustly documented a profound sialyl linkage switching.

Facilitated by their exposed position and flexible nature, the negatively charged sialic acids in the glycocalyx are important molecular players in a broad range of biological processes in the extracellular milieu. For example, sialylation is known to play crucial roles in immune modulation by influencing cell-cell interactions, receptor signaling and cytokine secretion [62, 64, 90]. In line with the observed shift towards α2,6-sialylation in anti-inflammatory macrophages, α2,6-linked sialic acids were previously associated with immunosuppressive phenotypes in the TME in the context of cancer progression and metastasis [36, 61]. In fact, a lectin array-focused study of the human leukemia monocytic cell line (THP-1) previously reported an elevation of α2,6-sialylation in anti-inflammatory “macrophage-like” cells [39] while another THP-1-based study found altered (de)sialylation and changes in sialyltransferase (ST) expression upon polarization [40]. Our study documenting a profound sialyl linkage switching in human monocyte-derived (primary) macrophages is an important extension of these previous *in vitro* findings as immortalized THP-1 cells represent an artificial model system that therefore may not accurately reflect the dynamic and non-templated glycosylation processes occurring in macrophages *in vivo*.

No prominent sialylation changes were detected for the surface *O*-glycome suggesting, in line with literature [91], that sialylation of *N*- and *O*-glycans is independently regulated by a broad family of STs with different yet overlapping substrate specificity. While protein-linked *N*- and *O*-glycans are prominent molecular features of the macrophage glycocalyx, changes in terminal sialylation of *N*-glycans may arguably have a greater functional impact as the sialic acid residues of these more elongated and flexible carbohydrates are more exposed to the extracellular environment compared to those carried by the more truncated *O*-glycans closer to the base of the glycocalyx. In support, suppression of the *N*-glycan sialylation of the anti-inflammatory macrophages (*ST6GAL1* KO), unmasked α2,3-sialyl glycoepitopes carried by the underlying *O*-glycans as measured by a previously unappreciated MAL-I recognition of WT cells using lectin fluorescence microscopy (Figure 4F). A limitation of this study is that we did not survey the sialylation of glycosphingolipids (e.g. gangliosides) in the macrophage glycocalyx, which therefore with the exception of a THP-1-focused study showing GM3 ganglioside elevation upon monocyte differentiation [42] remains largely undefined.

Importantly, our study found that morphologically heterogenous nanotubes carrying exposed α2,6-sialylation protrude in highly branched yet well-coordinated networks from and often between anti-inflammatory macrophages, cellular structures that were strikingly absent in pro-inflammatory macrophages (Figure 3B). To our knowledge, only scattered reports of nanotubes (alternatively referred to as nanotunnels, membrane nanotubes, tunnelling nanotubes [TNTs] or cytonemes) are available in the scientific literature with some reports in astrocytes [68] and cancer cells [69]. Relevant to our observation, a breast cancer-focused study performed in murine and zebrafish model systems found that TNTs between macrophages and tumor cells promote invasion and metastasis through epidermal growth factor (EGF) - EGF receptor signaling pathways with TNF-α induced protein 2 (TNFAIP2) found to be essential for TNT formation. [70]. The same group also reported that actin polymerization is essential for TNT formation and stability in murine RAW/LR5 monocyte/macrophage-like cells and identified that Rho GTPase effectors are important in TNT biogenesis in those cell systems [92]. Contrasting this and other mouse studies suggesting that actin form a key filamentous component of cell protrusions [93, 94], we did not identify actin to be a major nanotube component in the investigated primary anti-inflammatory macrophages. Together with the fact that we observed TNFAIP2 elevated in pro-rather than anti-inflammatory macrophages (Supplementary Table S6) indicates that the α2,6-sialylated nanotubes in our study may be different from the previously reported TNTs in non-human model systems. Sialobiology may be particularly sensitive to species variations given the absence of *N*-glycolylneuraminic acid in humans while such sialic acid species are dominant features in murine and other non-human mammalian model systems. Our study therefore offers new insights into the genesis, molecular makeup and functions of nanotubes in human anti-inflammatory macrophages through the lens of glycobiology.

Future efforts are warranted to determine the identity of the nanotubular proteins and their glycosylation sites as well as the biomolecular content and architecture of the nanotubes e.g. lipid membrane enclosed or non-membranous “naked” filamentous protein strings. A study performed in human monocyte-derived (primary) dendritic cells reported that nanotubes are capable of transferring calcium and other small molecules between cells upon mechanical stimulation [71] suggesting similar functions in the closely related anti-inflammatory macrophages, aspects we are currently exploring. We speculate that previous microscopy focused studies of human anti-inflammatory macrophages may have overlooked the nanotubular structures as they appeared clearly visible only upon SNA-based fluorescence microscopy. Even when probed with SNA, THP-1 cells polarized into anti-inflammatory “macrophage-like” cells appeared to lack nanotubes demonstrating the importance of avoiding this model system when studying macrophage glycobiology [40].

Accompanying macrophage polarization was a prominent remodeling of sialic acid-relevant enzymes and receptors providing clues to the biosynthetic mechanisms driving and the functional consequences resulting from the polarization-induced sialyl linkage switching. ST6GAL1, a prominent ST known to catalyze α2,6-sialylation of *N*-glycans [72, 74], was dramatically elevated in anti-inflammatory compared to pro-inflammatory macrophages suggesting the involvement of this glycoenzyme in the sialyl linkage switching of polarizing macrophages. Silencing of *ST6GAL1* in anti-inflammatory macrophages indeed reduced the SNA-reactivity and led to truncation and disintegration of the α2,6-sialylated nanotubes (Figure 4F) suggesting not only a biosynthetic, but also a functional involvement of ST6GAL1 in the glycophenotypic switching of polarizing macrophages.

ST6GAL1 has frequently been associated with cancer-promoting roles in the TME [34, 95]. For example, ST6GAL1-mediated α2,6-sialylation of tumor necrosis factor receptor 1 was reported to inhibit cancer cell apoptosis and facilitate formation of distant tumor sites [96]. In glioblastoma, ST6GAL1 also exhibited pro-tumorigenic roles by mediating sialylation of key surface receptors, such as integrins, critical for cell adhesion, migration and interaction [97]. Studies of THP-1 and RAW264.7 cells have associated ST6GAL1 and other STs with changes in sialylation during macrophage differentiation and polarization [39, 40]. Our study confirms the involvement of ST6GAL1 in the sialyl linkage switching upon polarization of human macrophages and expands on these findings by documenting new cancer-promoting functions of ST6GAL1 and α2,6-sialylation upon macrophage polarization. In addition to nanotube destruction, ST6GAL1 disruption led to impaired cellular motility, reduced macrophage-macrophage (homotypic) and macrophage-CRC cell (heterotypic) interactions and attenuated CRC proliferation compared to untreated (WT) anti-inflammatory cells. This demonstrates that ST6GAL1 not only catalyzes α2,6-sialylation critical for nanotube integrity and macrophage motility and interactions, but also serves as a pro-tumorigenic mediator between anti-inflammatory TAMs and CRC cells.

TAMs sense and respond to local changes in sialylation through sialic acid-binding immunoglobulin-like lectins (Siglecs), a broad family of signaling receptors that decorate macrophages and other tumor-infiltrating immune cell types. Our proteomic profiling revealed that Siglec-7, an inhibitory (ITIM-containing) receptor known to recognize α2,3/6/8-linked sialic acids on various biomolecules (gangliosides > *O*-glycoproteins > *N*-glycoproteins) [91, 98–100], is dramatically elevated in anti-inflammatory relative to pro-inflammatory macrophages suggesting its involvement in macrophage polarization or downstream functional events. In support, a transcriptomics study showed that Siglec-7 expression correlates with the infiltration rate of anti-inflammatory macrophages into glioma tissue and patient survival [101]. In addition, a recent study performed in mice found that Siglec-E (the mouse ortholog of human Siglec-7) is a major cancer-promoting TAM receptor responsive to tumor hyper-sialylation and reported that the efficacy of immune check point blockade can be improved by therapeutic desialylation and TAM repolarization [45]. Our observations of secreted Siglec-7 suggest extracellular roles beyond cell surface interactions in line with the extracellular role of secreted ST6GAL1 in cancer cell growth [102], but given the preference of Siglec-7 for ganglioside ligands (over *N*-glycans) it remains unknown if the polarization-induced sialyl linkage switching within the surface *N*-glycome is linked to Siglec-7 effector functions. While Siglec-9, which preferentially recognizes α2,3-sialylated *N*-glycans [103] and previously associated with TAM differentiation [104] and CRC prognosis and outcome [105], was found on both macrophage sub-population, both Siglec-1 (sialoadhesin, recognizing α2,3-sialyl epitopes) and Siglec-10 (recognizing α2,3/6-sialyl epitopes) [98, 103, 106, 107] were over-expressed in pro-relative to anti-inflammatory macrophages. Supporting their role in pro-inflammatory macrophages, Siglec-1 and -10 were previously found to exhibit anti-tumor activities including suppression of tumor growth [107–109]. Our study associates multiple Siglecs with the glycophenotypic modulation of TAM polarization, but further work is required to detail if, how and when Siglecs mechanistically tie into the sialyl linkage switching and effector functions of polarizing macrophages.

The anti-cancer effects observed upon ST6GAL1 inhibition in anti-inflammatory macrophages (reduced CRC cell adhesion and attenuated proliferation), raise the potential for therapeutic applications. Given their central role in cancer and other immune-related diseases, intense research efforts have been invested in identifying inhibitors against cancer-promoting STs [32, 110]. For example, a fluorinated sialic acid analogue (P-3Fax-Neu5Ac) was found to potently reduce α2,3/6-sialylation and lower the tumorigenic properties *in vitro* and *in vivo* in melanoma [111]. While reaching selectively and potency for individual STs have proven challenging, a recent lithocholic acid derivative FCW393 showed effective and selective inhibition of ST6GAL1 (IC_50_ 7.8 μM) and ST3GAL3 (9.45 μM) leading to reduced tumorigenic traits both *in vitro* and *in vivo* in breast cancer [112]. Given the increasing availability of effective ST6GAL1 inhibitors, studies are now underway in attempts to reduce in a controlled manner anti-inflammatory TAMs in the TME, with the aim to restore immune balance and favor tumor-fighting properties of the immune infiltrate. Relevant to our work, recent studies have shown that de-sialylation can disrupt polarization and shift TAMs towards a pro-inflammatory state [45, 104] demonstrating that repolarization is achievable through sialic acid modulation.

In this study, we report on the mechanistic basis for and functional consequences of the profound glycocalyx remodeling accompanying TAM polarization. The *ex vivo* approach allowed us to map with unprecedented depth and precision the glycophenotypic changes underpinning the polarization of human macrophages using powerful system glycobiology tools that have only recently become available [113], but it is acknowledged that this setup does not fully mimic the true cellular complexity of the local tissue environment in which macrophages undergo polarization. While the sialyl linkage switching was found to be reflected *in situ* in tumor tissues resected from CRC patients, future glycobiology-focused studies are required to expand on these findings to document the role of the polarization-induced sialyl linkage switching in TAMs and demonstrate *in vivo* the anti-tumor potential of ST6GAL1 inhibition. Conclusively, our study provides mechanistic insights into the glycobiology of TAMs, in turn, elevating our understanding of the TME and opening new avenues for advanced immuno-therapies against cancer.

## Data availability

The glycomics LC-MS/MS raw data were deposited to GlycoPOST [114] with the identifier GPST000502 (https://glycopost.glycosmos.org/preview/93266032467326f9b747f9, pin code: 6382). Proteomics LC-MS/MS raw data were deposited to the ProteomeXchange Consortium via PRIDE [115] partner repository with the identifier PXD057634 (Username, Password: J9R3SlHdJ3Gz) (see **Supplementary Table S7** for overview of data file).

## Author contributions

PD and MTA designed experiments. PD, NB, ZSB, AC, DK, MA, RK conducted experiments. PD, AC, JU, THC, RK, MTA analyzed data. BT and MTA provided supervision and resources. PD and MTA wrote manuscript. All authors have edited and read the manuscript.

## Acknowledgements (coauthors to confirm or add details)

We acknowledge the Australian Proteome Analysis Facility and the Microscopy Unit, Faculty of Science and Engineering, Macquarie University for their support and assistance in this work. PD was supported by International Macquarie University Research Excellence Scholarship and a Top-up scholarship by Hebrew University. NB was supported by a Macquarie University Research Excellence Scholarship. AC is supported by a Research Training Program scholarship funded by the Australian Government. THC is supported by an International Research Training Program Scholarship. DK and MA were supported by the Australian Government via the National Health and Medical Research Council (NHMRC) Ideas grant (GNT2018947). RK is supported by KAKENHI (24K17793 & 23K19347). MTA is the recipient of an ARC Future Fellowship (FT210100455).

## Supplemental information

Supplementary data accompany this paper. Supplementary Figures (Supplementary Figures S1-S12) are provided in a separate PDF document, Supplementary Tables (Supplementary Table S1-S6) are compiled in Microsoft Excel document and Supplementary Videos (Supplementary Videos S1-S3) are provided as Microsoft PPT files.

**Supplementary Figure S1.** Monocyte isolation, differentiation and validation.

**Supplementary Figure S2.** Sialyl linkage switching on *N*-glycans in the cell lysate fraction of pro- and anti-inflammatory macrophages.

**Supplementary Figure S3.** Validation of cell surface proteins in the biotinylated fraction of pro- and anti-inflammatory macrophages.

**Supplementary Figure S4.** Opposite (but modest) sialyl linkage switching of a surface *O*-glycan upon macrophage polarization.

**Supplementary Figure S5.** Distinct SNA and MAL-I reactivity in pro- and anti-inflammatory macrophages.

**Supplementary Figure S6.** Nanotubes decorated with α2,6-sialylation protrude from anti-inflammatory macrophages.

**Supplementary Figure S7**. Actin-less nanotubes in anti-inflammatory macrophages.

**Supplementary Figure S8.** ST6GAL1 depletion impairs nanotube-mediated heterotypic interactions between anti-inflammatory macrophages and CRC cells.

**Supplementary Figure S9**. *ST6GAL1* silencing in anti-inflammatory macrophages reduces CRC cell proliferation.

**Supplementary Figure S10.** Proteome remodeling in pro- and anti-inflammatory macrophages.

**Supplementary Figure S11.** Siglec expression in pro- and anti-inflammatory macrophages.

**Supplementary Figure S12.** Absence of TAMs and sialyl linkage switching in normal adjacent colon tissue from CRC patients.

**Supplementary Table S1**. *N*-glycans in cell lysate of pro- and anti-inflammatory macrophages.

**Supplementary Table S2**. Proteins identified in the biotinylated and non-biotinylated fractions of pro- and anti-inflammatory macrophages.

**Supplementary Table S3**. DAVID pathway enrichment analysis of the biotinylated and non-biotinylated fraction of pro- and anti-inflammatory macrophages.

**Supplementary Table S4**. Sialyl linkage profiling of abundant sialo-N-glycans identified on the cell surface of pro- and anti-inflammatory macrophages.

**Supplementary Table S5**. Sialyl linkage profiling of abundant sialo-O-glycans identified on the cell surface of pro- and anti-inflammatory macrophages.

**Supplementary Table S6**. Proteomics of the cell lysate and surface fractions of pro- and anti-inflammatory macrophages.

**Supplementary Table S7**. Overview of deposited raw LC-MS/MS data files.

**Supplementary Video S1.** ST6GAL1 disruption leads to nanotube disintegration, and reduced motility and homotypic interactions of anti-inflammatory macrophages.

**Supplementary Video S2.** ST6GAL1 disruption reduces motility of anti-inflammatory macrophages and attenuates their heterotypic interactions with CRC cells.

**Supplementary Video S3.** ST6GAL1 disruption in anti-inflammatory macrophages impairs nanotube-mediated heterotypic interactions to CRC cells.

**Supplementary Video S1. ST6GAL1 disruption leads to nanotube disintegration, and reduced motility and homotypic interactions of anti-inflammatory macrophages.** Live cell recordings of WT (left) and *ST6GAL1* KD (right) anti-inflammatory macrophages were performed for 21 h. Different colored arrows indicate macrophage-macrophage interactions. Insert: Migration was tracked for the individual macrophages (in different colors) over the 21 h period. Representative videos shown from separate live recordings of biological duplicates. Scale bar = 50 µm.

**Supplementary Video S2. ST6GAL1 disruption reduces motility of anti-inflammatory macrophages and attenuates their heterotypic interactions with CRC cells.** Live cell recordings of WT (left) and *ST6GAL1* KD (right) anti-inflammatory macrophages were performed in co-cultures with LIM2405 CRC cells for 29 h. Colored triangles indicate macrophage-CRC cell interactions while colored arrows indicate CRC cells detaching from the substrate (loss of cell adhesion) or being phagocytosed. Representative videos shown from separate live recordings of three biological replicates (n = 3). Scale bar = 50 µm.

**Supplementary Video S3. ST6GAL1 disruption in anti-inflammatory macrophages impairs nanotube-mediated heterotypic interactions to CRC cells**. Live cell recording of untreated (WT) or *ST6GAL1* silenced anti-inflammatory macrophages in co-cultures with LIM2405 CRC cells for 29 h. Colored triangles indicate intact or fragmented/truncated thin nanotubes that enable or prevent heterotypic macrophage-CRC interactions in the WT or ST6GAL1-disrupted anti-inflammatory macrophages. Colored arrows indicate CRC cell proliferation. Representative video shown from separate live recordings of three biological replicates (n = 3). Scale bar: 50 µm.

