## Supplementary Figure S1-S12 for "ST6GAL1-mediated sialyl linkage switching in tumor-associated macrophages drives cancer-promoting nanotubes carrying α2,6-sialylation in anti-inflammatory cells"

### for


Running title: ST6GAL1 drives cancer-promoting α2,6-sialylated nanotubes in anti-inflammatory TAMs


Keywords: ST6GAL1, sialylation, macrophage, nanotubes, polarization, sialyltransferase, Siglecs, colorectal cancer, interactions, proliferation.



Corresponding author
Associate Professor Morten Thaysen-Andersen
School of Natural Sciences
Macquarie University
Sydney, Australia

ORCID: 0000-0001-8327-6843

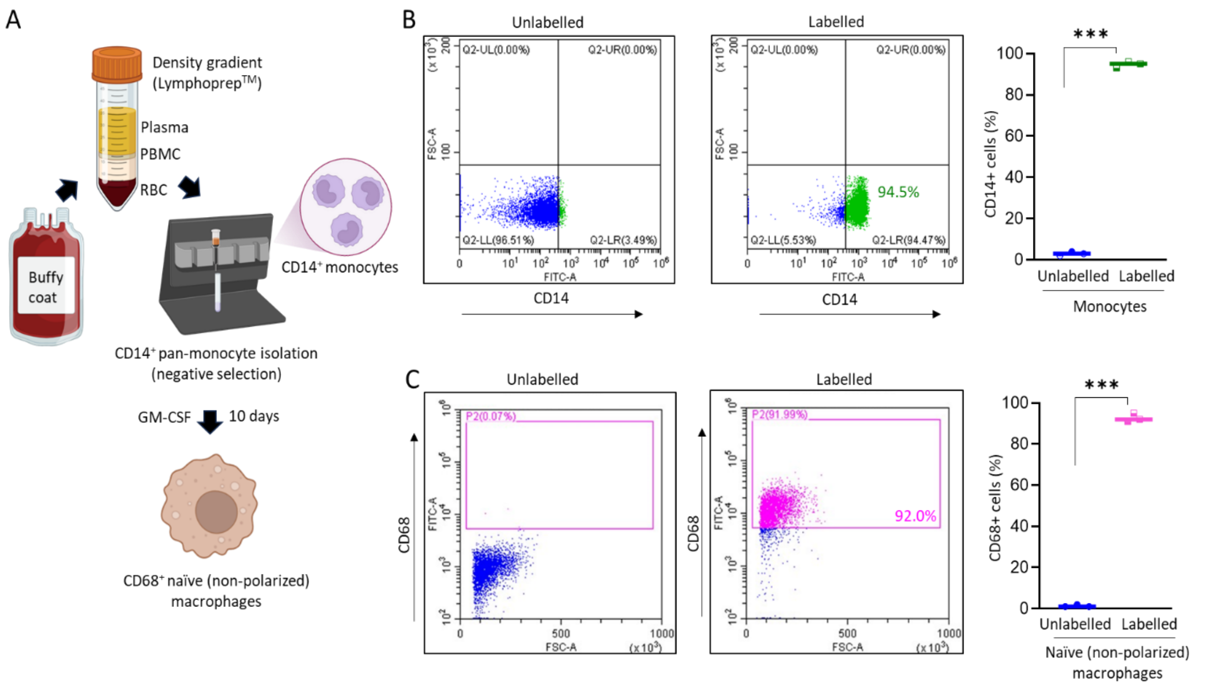


**Supplementary Figure S1. Monocyte isolation, differentiation and validation. A)** Schematic representation of CD14+ monocyte isolation from buffy coat of healthy donors (n = 4). Buffy coats were processed using density gradient centrifugation with Lymphoprep to obtain a PBMC layer that was negatively selected for CD14+ monocytes prior to GM-CSF-induced differentiation into naïve (non-polarized) macrophages over 10 days. Flow cytometry-based validation of the **B**) monocytes (CD14+) and **C**) naïve macrophages (CD68+) against unlabeled populations (autofluorescence controls). Two-tailed paired t-test (***$p < 0.001$, n = 3).

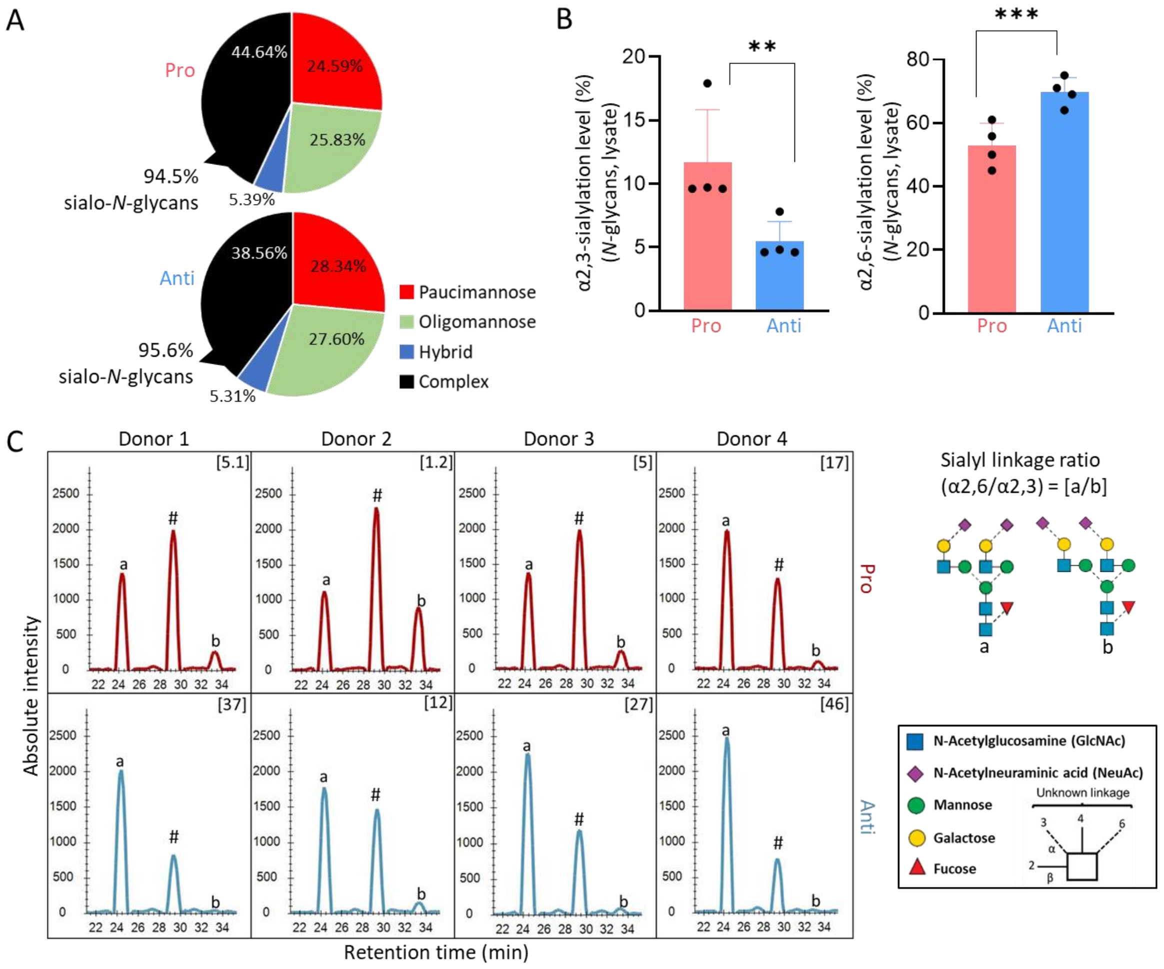


**Supplementary Figure S2. Sialyl linkage switching on *N*-glycans in the cell lysate fraction of pro- and anti-inflammatory macrophages. A**) *N*-glycan type distribution and degree of sialylation of the complex type *N*-glycans in the cell lysate fraction from pro- (top) and anti- (bottom) inflammatory macrophages using PGC-LC-MS/MS glycomics data. See key for glycan types. **B**) Global α2,3- (left) and α2,6- (right) sialylation levels based on PGC-LC-MS/MS *N*-glycomics data of cell lysate fractions, see Supplementary Table S1 for tabulated glycomics data. Data are presented as mean ± SD, and individual data points are shown. Two-tailed paired t-test (**$p < 0.01$, n = 4). **C**) Extracted ion chromatograms of three discrete sialo-isomers of a prominent biantennary disialylated *N*-glycan (*m/z* 1184.4) carrying both α2,6- (a isomer), α2,3- (b isomer) and a combination of both α2,6/α2,3- (marked with #) sialylation in the cell lysate fraction from pro- (top) and anti- (bottom) inflammatory macrophages using PGC-LC-MS/MS glycomics data. The relative levels of only the a and b isomers were used to establish the α2,6/α2,3-sialyl linkage ratio (see inserts in each chromatogram). Glycans are depicted according to SNFG nomenclature [1].

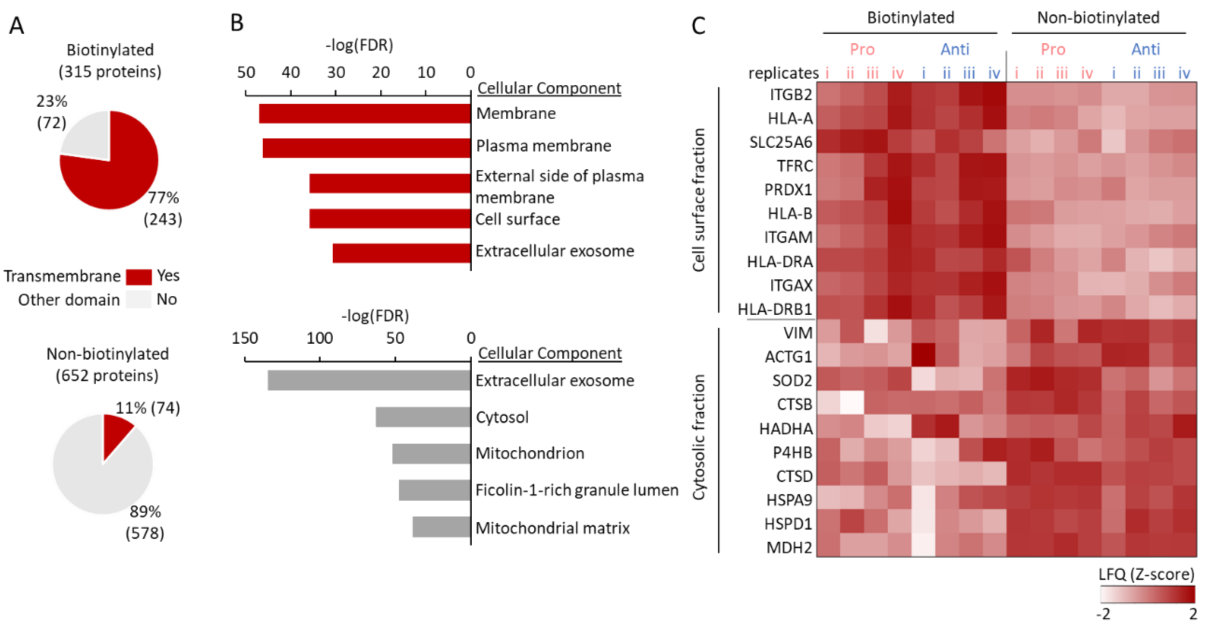


**Supplementary Figure S3. Validation of cell surface proteins in the biotinylated fraction of pro- and anti-inflammatory macrophages.** **A**) Proteins identified in the biotinylated (top) and non-biotinylated (bottom) fractions were mapped for the presence of transmembrane domains. **B**) Enrichment of cellular components (Gene Ontology) in the biotinylated (top) and non-biotinylated (bottom) fractions. **C**) The ten most differentially abundant proteins from each fraction are displayed as a heat map using log-transformed label-free quantified (LFQ) Z-scores. Cellular component enrichment analysis was performed using Database for Annotation, Visualization and Integrated Discovery (DAVID) Bioinformatics (NIH) with corrected $p < 0.05$ as significance threshold. Transmembrane domain annotations were performed using the UniProt ID mapping tool.

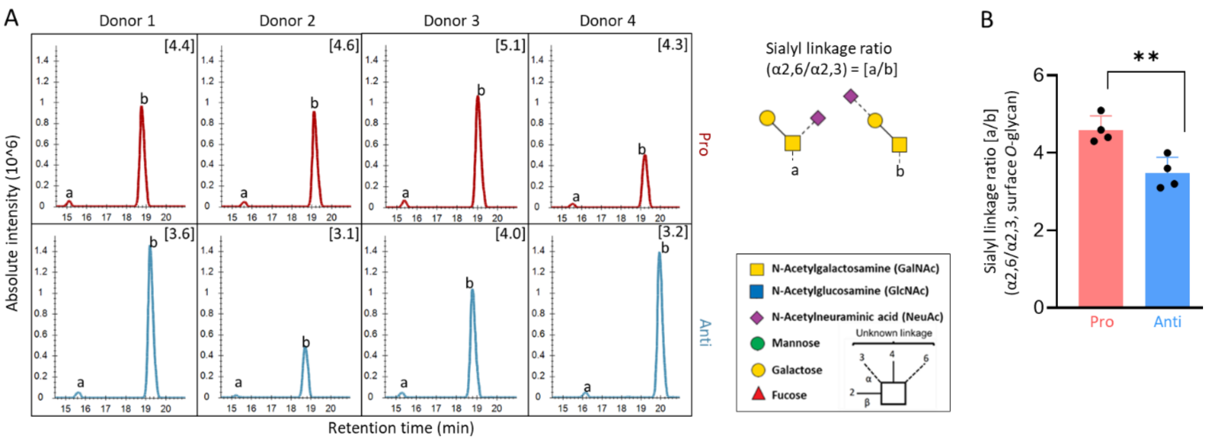


**Supplementary Figure S4. Opposite (but modest) sialyl linkage switching of a surface *O*-glycan upon macrophage polarization. A**) Extracted ion chromatogram of two discrete sialo-isomers of a prominent trisaccharide *O*-glycan (*m/z* 675.3) ($Hex_1HexNAc_1NeuAc_1$) carrying both α2,6- (a isomer) and α2,3- (b isomer, see insert for structures) sialylation in the cell surface fraction from pro- (top) and anti- (bottom) inflammatory macrophages using PGC-LC-MS/MS glycomics data. The relative levels of the a and b isomers were used to establish the α2,6/α2,3-sialyl linkage ratio (see inserts in XIC plots). Glycans are depicted according to SNFG nomenclature [1]. **B**) The α2,6/α2,3-sialyl linkage ratios [a/b] of this specific surface *O*-glycan were compared between the pro- and anti-inflammatory macrophages. Data are presented as mean ± SD, and individual data points are shown. Paired two-tailed t-test (**$p < 0.01$).

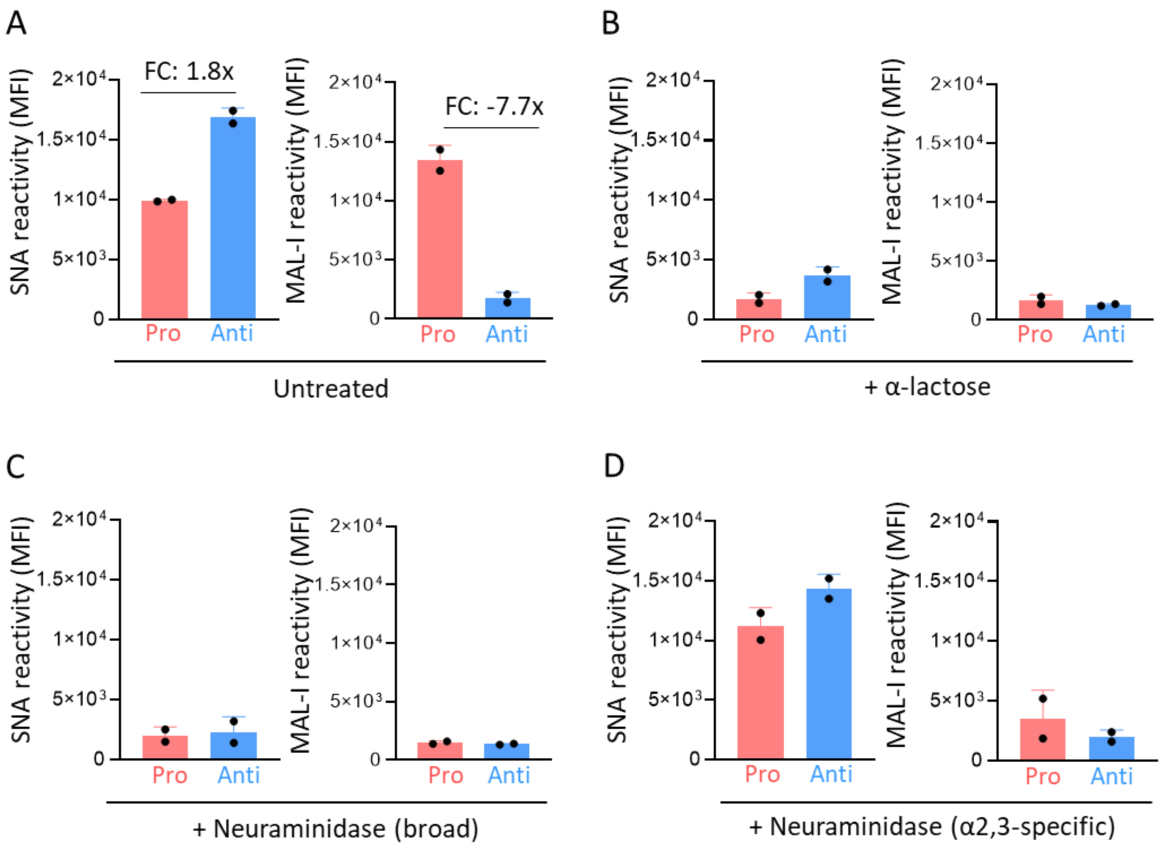


**Supplementary Figure S5. Distinct SNA and MAL-I reactivity in pro- and anti-inflammatory macrophages. A**) Lectin flow cytometry data of untreated pro- and anti-inflammatory macrophages (see Figure 2D) were quantified based on the mean fluorescence intensity (MFI) of PE (conjugated with SNA, left graph) and FITC (conjugated with MAL-I, right graph) gated channels. Several controls were carried out to ensure lectin recognition specificity including pre-treatment of lectins with **B**) α-lactose (see Figure 2E for flow cytometry data) and pre-treatment of cells with **C**) broad neuraminidase (see Figure 2F for flow cytometry data) and **D**) α2,3-specific neuraminidase (see Figure 2G for flow cytometry data). For B-D, MFI was plotted as in A. For all graphs, data are presented as mean ± SD, and individual data points are shown. FC: Fold-change from biological duplicates (n = 2).

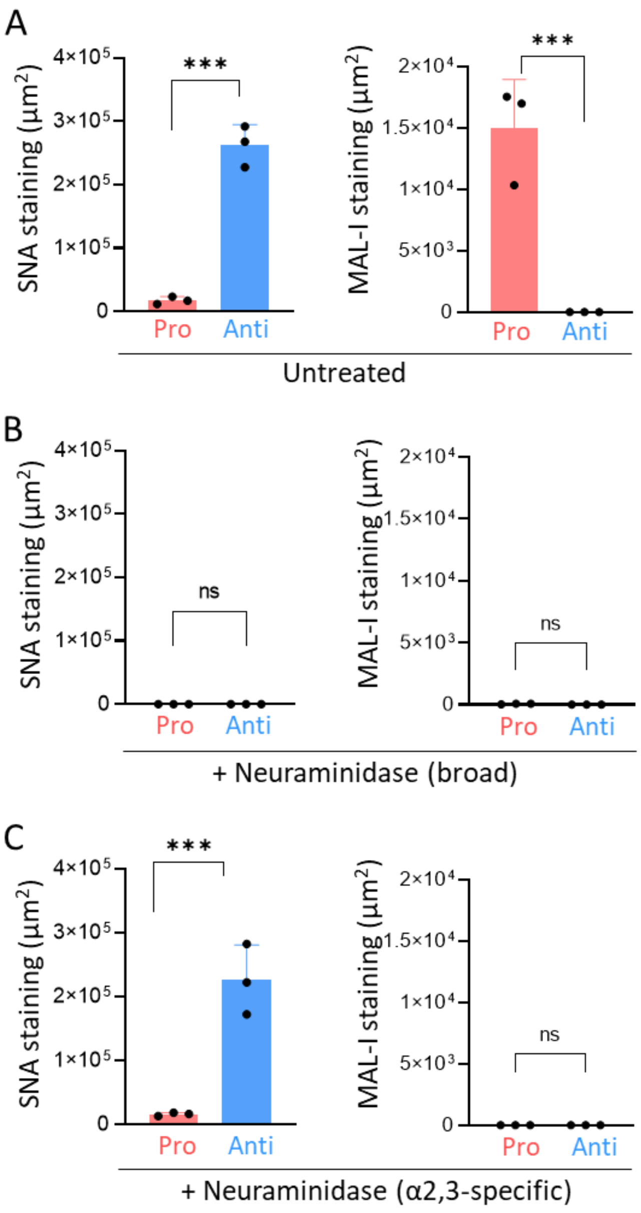


**Supplementary Figure S6. Nanotubes decorated with α2,6-sialylation protrude from anti-inflammatory macrophages.** Lectin fluorescence-based quantitation of SNA (left) and MAL-I (right) reactivity measured as stained area per microscopic field ($\mu m^2$) of **A**) untreated pro- and anti-inflammatory macrophages and of cells pre-treated with **B**) neuraminidase (broad) and **C**) α2,3-specific neuraminidase. See Figure 3 for immune fluorescence microscopy. Data are presented as mean ± SD, and individual data points are shown. Two-tailed paired t-test (***$p < 0.001$, ns, non-significant $p \geq 0.05$, n = 3).

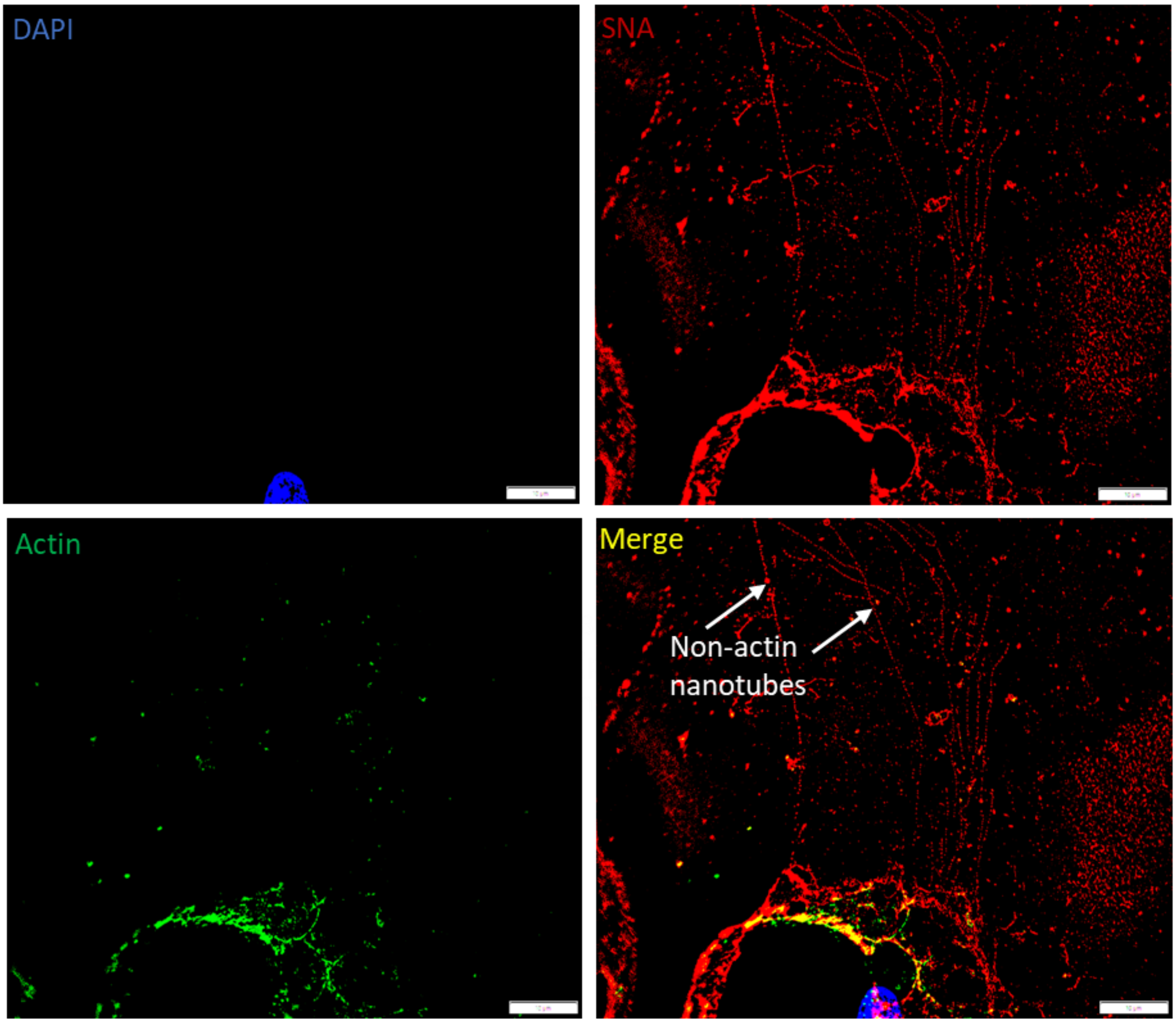


**Supplementary Figure S7**. **Actin-less nanotubes in anti-inflammatory macrophages.** Fluorescence microscopy of anti-inflammatory macrophages stained with DAPI (blue), SNA (red), and ActinGreen488 ReadyProbe (green). The merged image was generated by combining all three channels. White arrows: Examples of SNA-reactive nanotubes devoid of exposed actin. Scale bar: 10 µm.

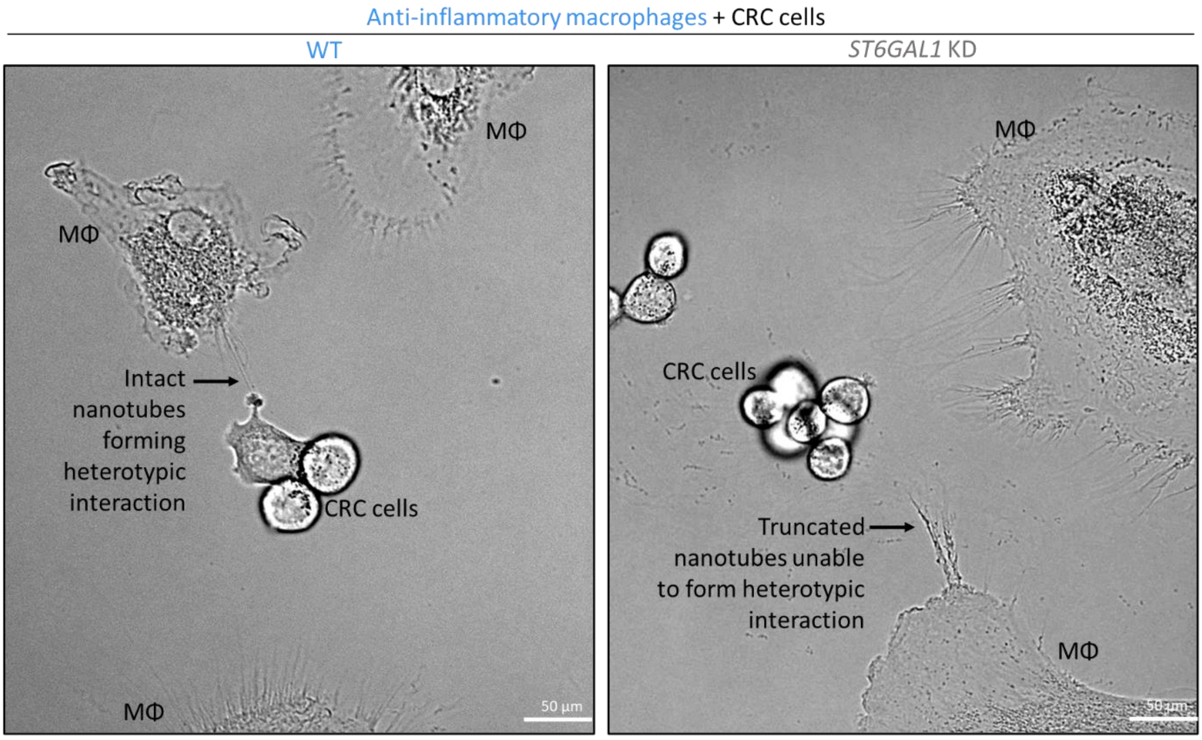


**Supplementary Figure S8. ST6GAL1 depletion impairs nanotube-mediated heterotypic interactions between anti-inflammatory macrophages and CRC cells**. Anti-inflammatory macrophages (WT, left and *ST6GAL1* KD, right) were co-cultured with CRC cells (LIM2405, labelled) for 72 h and their heterotypic interactions then monitored over a 29-h period using live cell recordings. Representative images (of three independent live cell recording experiments) showing nanotube-mediated heterotypic interactions to the CRC cells (WT) and truncated nanotubes unable to form connections to the CRC cells (*ST6GAL1* KD) were captured from the live cell recordings, see **Supplementary Video S3**. Scale bars: 50 µm.

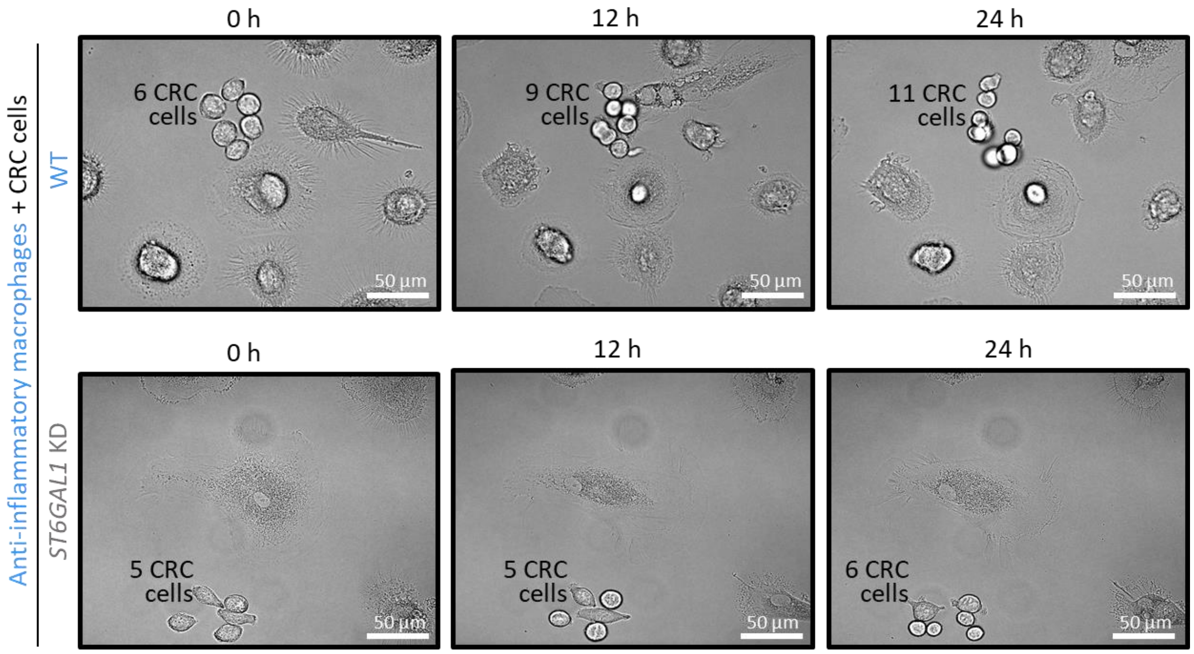


**Supplementary Figure S9**. ***ST6GAL1* silencing in anti-inflammatory macrophages reduces CRC cell proliferation**. Anti-inflammatory macrophages (WT, top row and *ST6GAL1* KD, bottom row) were co-cultured with CRC cells (LIM2405, labelled) for 72 h, after which the CRC cells were monitored for cell proliferation over a 29-h period using live cell recording. CRC cell counts are indicated at 0 h (as starting time point), 12 h and 24 h. Presentative images from three independent experiments (n = 3) were captured from the live cell recordings, see **Supplementary Video S2**. Scale bars: 50 µm.

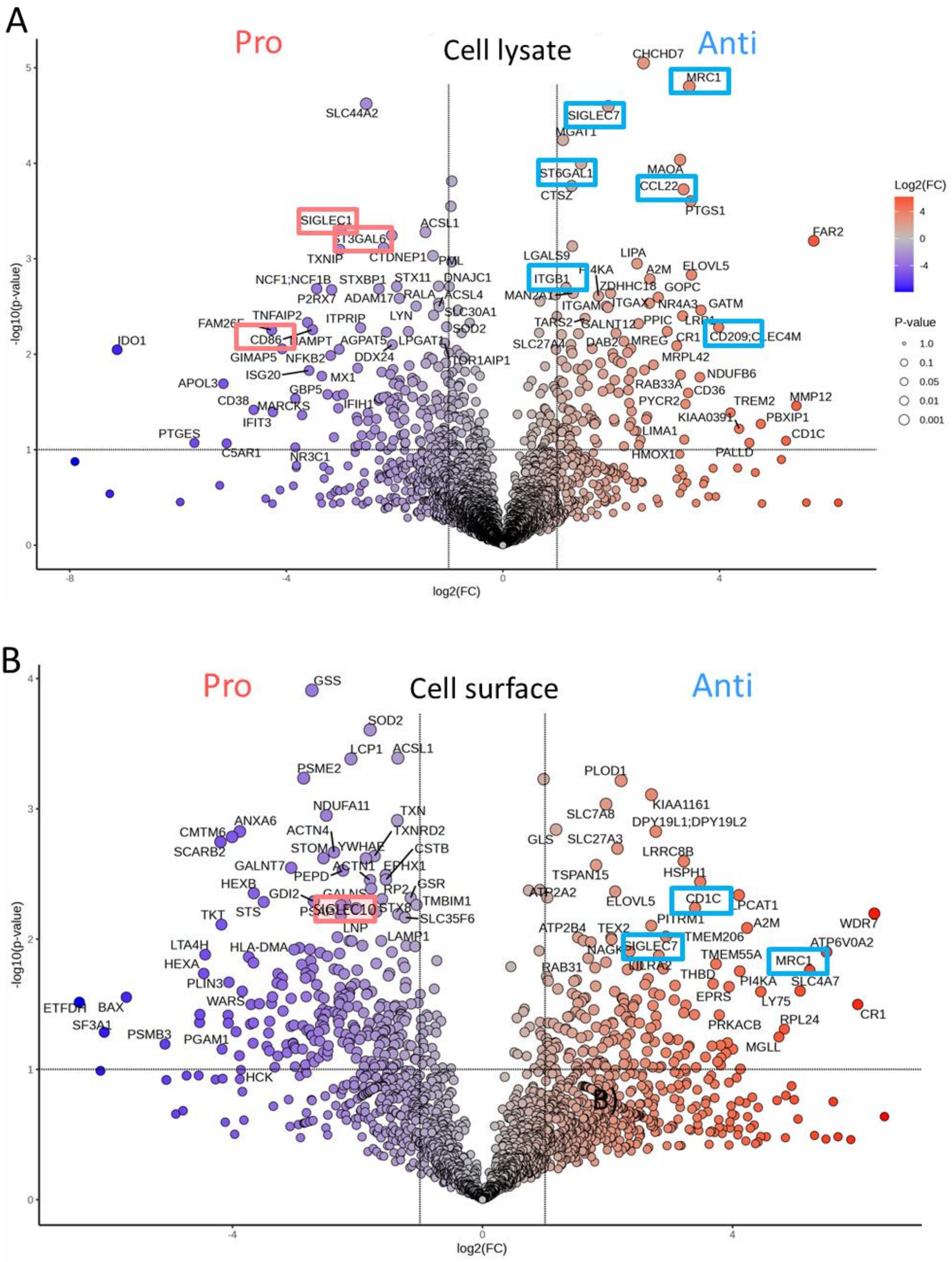


**Supplementary Figure S10. Proteome remodeling in pro- and anti-inflammatory macrophages.** Protein levels were quantified and compared in the **A**) cell lysate and **B**) cell surface fractions of pro- and anti-inflammatory macrophages using quantitative proteomics. x-axis: Log2 fold change (see insert for intensity scale). Sialic acid-related proteins and other key proteins elevated in pro-inflammatory (red boxes) and anti-inflammatory (blue boxes) macrophages are highlighted. The FC threshold for significance was set to 2 with $p < 0.05$. Paired t-tests (n = 4).

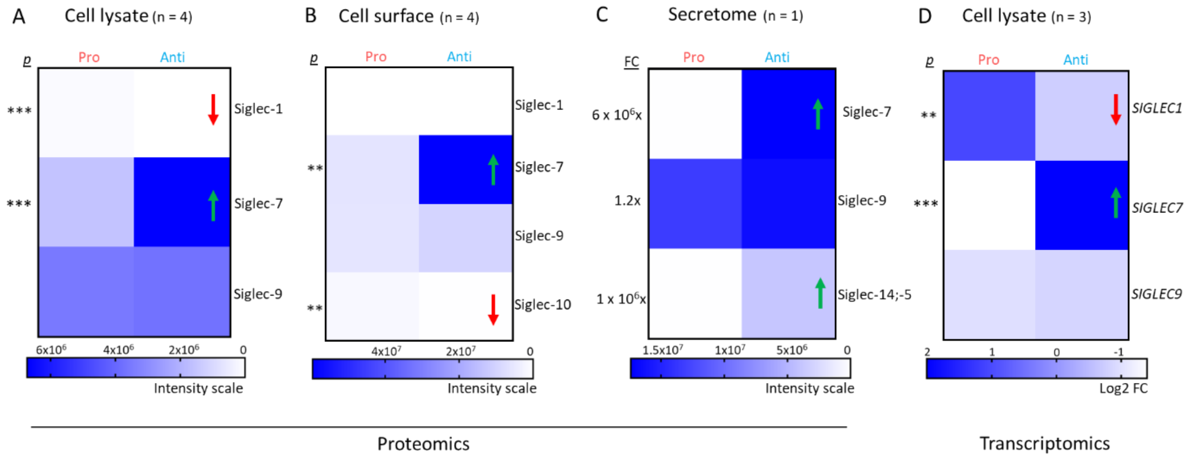


**Supplementary Figure S11. Siglec expression in pro- and anti-inflammatory macrophages.** Relative levels of sialic acid-binding immunoglobulin-like lectins (Siglecs) identified at the protein level in the **A**) cell lysate, **B**) cell surface and **C**) secretome using proteomics and **D**) at the gene expression level in the cell lysate using transcriptomics of pro- and anti-inflammatory macrophages. For both A-B, two-tailed, paired t-test, ***$p$ < 0.001, **$p$ < 0.01, *$p$ < 0.05 (n = 4). For C, FC: fold change (n = 1). See insert for intensity scale. Proteins/genes significantly higher (green arrow) or lower (red arrow) in anti-inflammatory compared to pro-inflammatory macrophages are indicated.

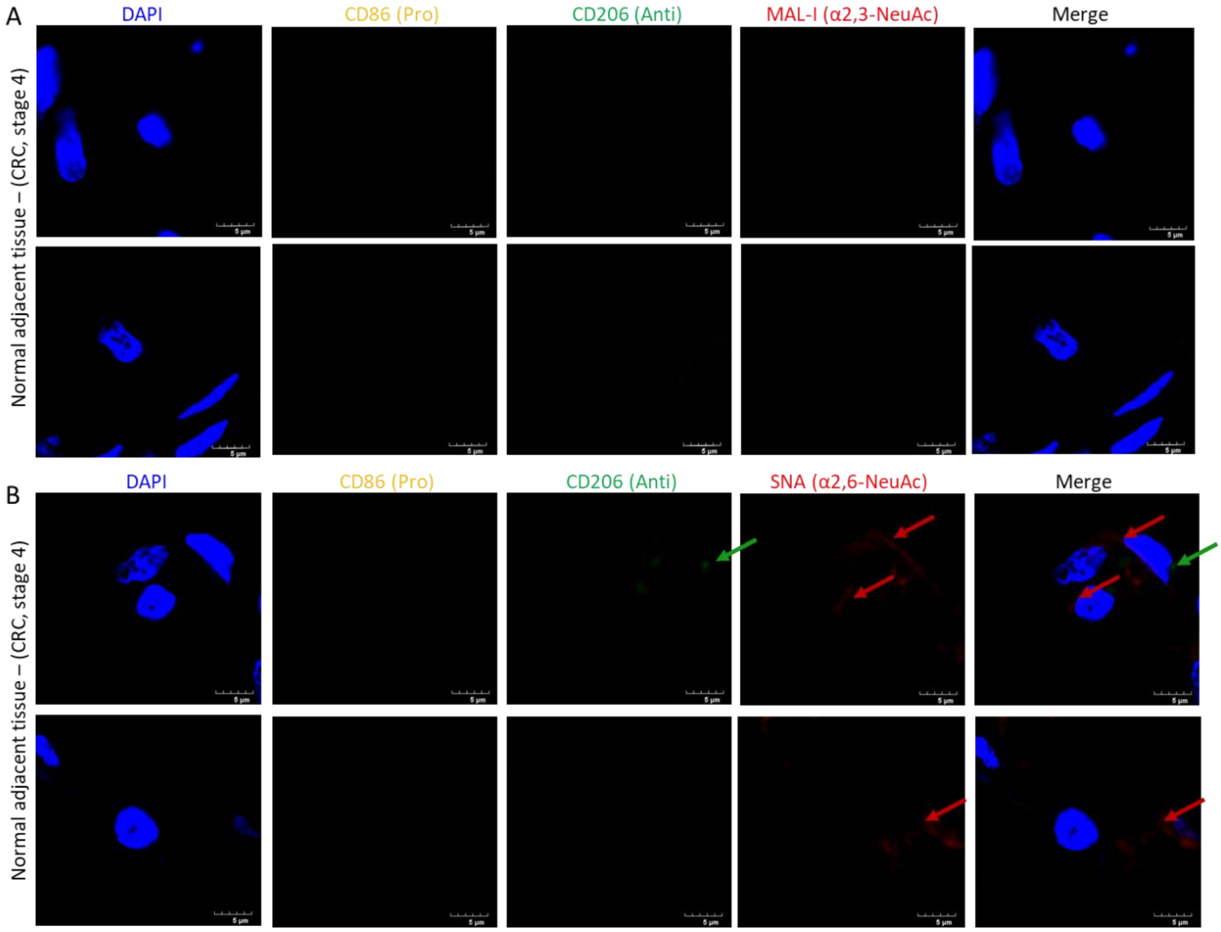


**Supplementary Figure S12. Absence of TAMs and sialyl linkage switching in normal adjacent colon tissue from CRC patients. A)** SNA- and **B**) MAL-I-based lectin histochemistry and immunohistochemistry (IHC) of normal adjacent tissue collected away from the tumor region of CRC patients with stage 4, see Figure 5 for lectin histochemistry and IHC of tumor tissues. Tissue sections were deparaffinized and stained with DAPI (nuclei), markers for pro-inflammatory (CD86, yellow) and anti-inflammatory (CD206, green) TAMs and sialyl-linkage recognizing lectins i.e. SNA (α2,6-sialic acid reactive) and MAL-I (α2,3-sialic acid reactive). Merged images were generated by combining all four channels. Arrows: Examples of unique features indicating a lack of co-localization. Scale bar: 5 µm. For both A-B, two representative images are shown from three different tissue sample slides (n = 3).

**References used in the supplementary material**

1. Neelamegham, S., et al., *Updates to the Symbol Nomenclature for Glycans guidelines.* Glycobiology, 2019. **29**(9): p. 620-624.
