## Supplementary figures and images for "ST6GAL1-mediated sialyl linkage switching in tumor-associated macrophages drives cancer-promoting nanotubes carrying α2,6-sialylation in anti-inflammatory cells"

### Supplementary Video S1

#### Slide 1
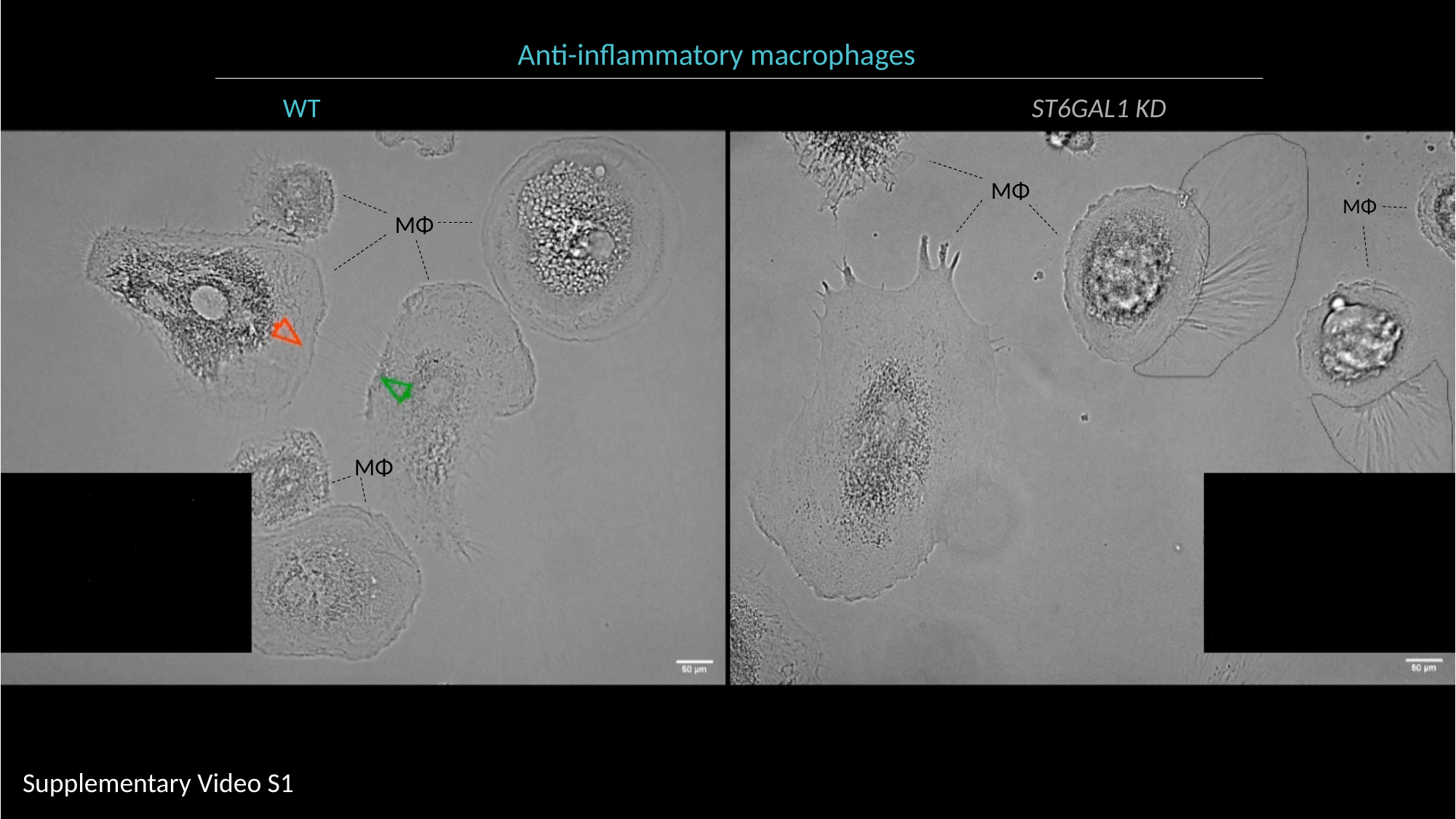

Anti-inflammatory macrophages
WT ST6GAL1 KD
MΦ
MΦ
MΦ
MΦ
Supplementary Video S1
