## Supplementary Video S2 for "ST6GAL1-mediated sialyl linkage switching in tumor-associated macrophages drives cancer-promoting nanotubes carrying α2,6-sialylation in anti-inflammatory cells"

### Slide 1
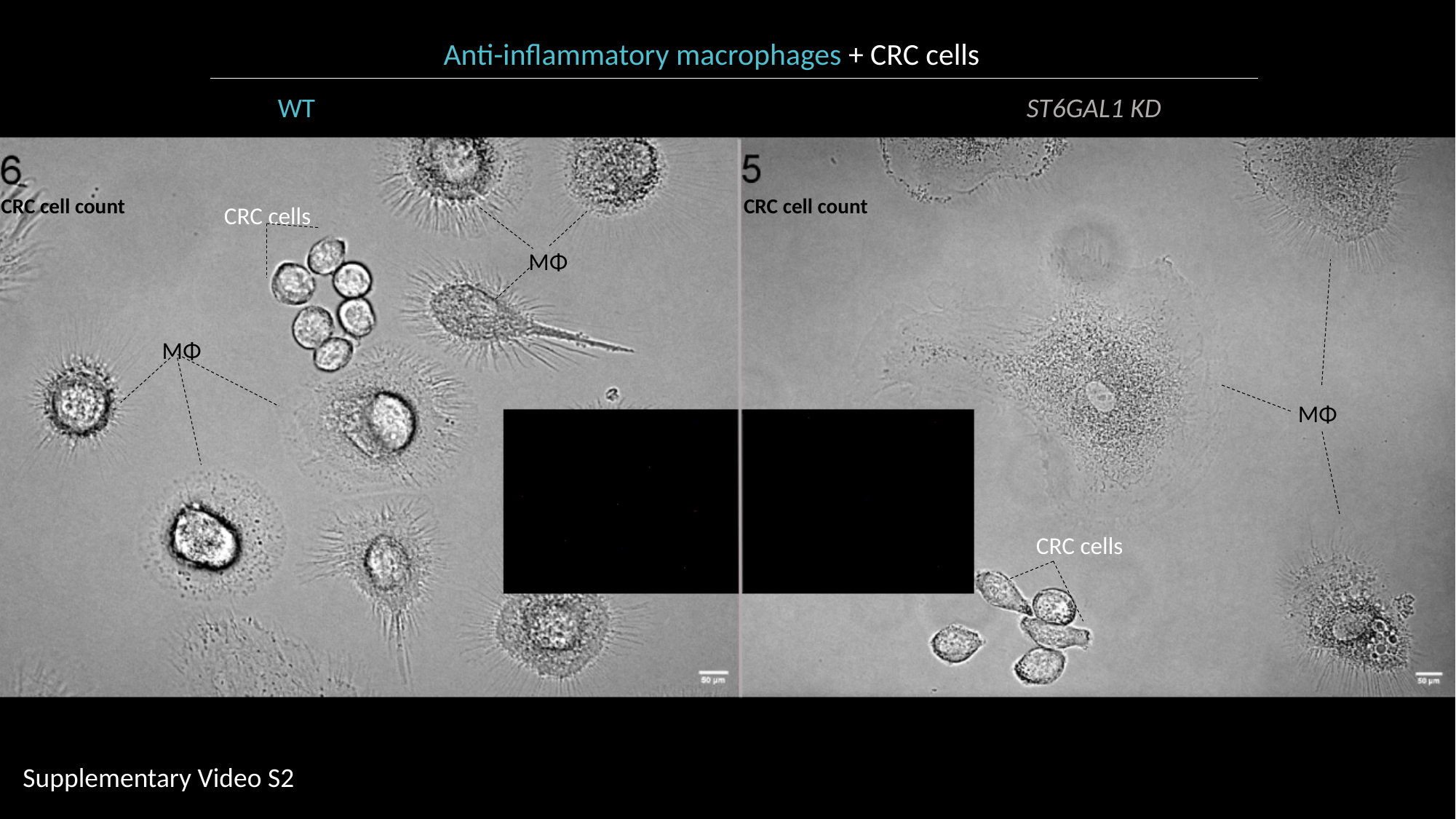

Anti-inflammatory macrophages + CRC cells
WT ST6GAL1 KD
CRC cell count
CRC cell count
CRC cells
MΦ
MΦ
MΦ
CRC cells
Supplementary Video S2
